# Supervised machine learning reveals introgressed loci in the genomes of *Drosophila simulans* and *D. sechellia*

**DOI:** 10.1101/170670

**Authors:** Daniel R. Schrider, Julien Ayroles, Daniel R. Matute, Andrew D. Kern

## Abstract

Hybridization and gene flow between species appears to be common. Even though it is clear that hybridization is widespread across all surveyed taxonomic groups, the magnitude and consequences of introgression are still largely unknown. Thus it is crucial to develop the statistical machinery required to uncover which genomic regions have recently acquired haplotypes via introgression from a sister population. We developed a novel machine learning framework, called FILET (Finding Introgressed Loci via Extra-Trees) capable of revealing genomic introgression with far greater power than competing methods. FILET works by combining information from a number of population genetic summary statistics, including several new statistics that we introduce, that capture patterns of variation across two populations. We show that FILET is able to identify loci that have experienced gene flow between related species with high accuracy, and in most situations can correctly infer which population was the donor and which was the recipient. Here we describe a data set of outbred diploid *Drosophila sechellia* genomes, and combine them with data from *D. simulans* to examine recent introgression between these species using FILET. Although we find that these populations may have split more recently than previously appreciated, FILET confirms that there has indeed been appreciable recent introgression (some of which might have been adaptive) between these species, and reveals that this gene flow is primarily in the direction of *D. simulans* to *D. sechellia*.

**AUTHOR SUMMARY:** Understanding the extent to which species or diverged populations hybridize in nature is crucially important if we are to understand the speciation process. Accordingly numerous research groups have developed methodology for finding the genetic evidence of such introgression. In this report we develop a supervised machine learning approach for uncovering loci which have introgressed across species boundaries. We show that our method, FILET, has greater accuracy and power than competing methods in discovering introgression, and in addition can detect the directionality associated with the gene flow between species. Using whole genome sequences from *Drosophila simulans* and *Drosophila sechellia* we show that FILET discovers quite extensive introgression between these species that has occurred mostly from *D. simulans* to *D. sechellia.* Our work highlights the complex process of speciation even within a well-studied system and points to the growing importance of supervised machine learning in population genetics.

## INTRODUCTION

Up to 10% of animal [1] and plant [2] species have the ability to hybridize with other species. Our recent ability to collect large-scale genomic data has confirmed that hybridization is common in nature. Indeed the ubiquity of hybridization upon secondary contact raises the question of how large a role hybridization plays in the emergence or collapse of new lineages [3].

Three general patterns have emerged from recent efforts to search for introgression in genomic data. First, whole-genome sequencing has shown that introgression occurs in all taxa for which its signature has been systematically sought (primates reviewed in [4], plants in [5, 6], fungi [7] and oomycetes in [8]). In general, genetic exchange between species through fertile hybrids might be common between closely related species [9–13] but can also occur between divergent species [14–17].

Second, introgression is heterogeneously distributed across the genome. For instance, mitochondrial genome exchange is surprisingly common (e.g., [18–20] among many) between species, whereas sex chromosomes are less likely to cross species-boundaries, perhaps due to their disproportionate role in hybrid incompatibilities [17, 21–24]. Generally it seems that functional regions of the genome might be less likely to participate in introgression. This is perhaps best known from the case of Neanderthal hybridization with non-African human populations [25, 26], which has left modern human genomes distinct gradients of introgression across different functional compartments of the genome.

Finally, the mode and intensity of natural selection acting on introgressed DNA can vary substantially. Loci that contribute to reproductive isolation, and as such to the persistence of species in the face of hybridization, should be less likely to be introgressed [27] as a result of purifying selection in hybrids. Additionally, introgressed haplotypes containing mildly deleterious variants may be purged after migrating into a population with a larger effective size where selection is more effective [28, 29]. On the other hand, much of the genome may be porous to introgression between closely related species if the net effect of such introgressed variation is fitness neutral. Of course genetic exchange between populations can also provide a source of adaptive alleles that may facilitate adaptation to new environments (reviewed in ref. [30]). Introgressions have indeed been shown to be involved in adaptation in animals (e.g. [31–33]), plants (e.g. [34]) and fungi [35]. For instance, adaptation to high altitude in Tibetans appears to have been caused by introgression of alleles from an archaic Denisovan-like source into modern humans [36]. Another particularly well-studied system of adaptive introgression comes from *Heliconius* butterflies where gene exchange has facilitated the origin and maintenance of mimetic rings [32] and even of hybrid species [37, 38].

Clearly, hybridization and introgression play an important role in shaping the landscape of genetic variation, thus if we wish to fully understand its evolutionary role a reliable framework for the inference of introgressed alleles is needed. Approaches to detect introgression in the genome fall into a few different camps. Genome-wide approaches can identify whether admixture has occurred in a set of populations. These include clustering methods which seek to infer which individuals are admixed and to assign a proportion of admixture to each individual without previous knowledge of the parental populations [39–41]. Some genome-wide approaches instead attempt to infer the directionality of introgression by examining allele frequency differences among populations [25, 42]. The main limitation of this class of methods is that they cannot identify which regions of the genome are likely to have crossed species boundaries.

On the other hand, locus-specific ancestry approaches (e.g. [43–47]) seek to uncover the mosaic of ancestry for each sampled haplotype, and thus can also identify portions of haplotypes that have been introgressed between species or populations. These fine-resolution approaches are powerful but often have assumptions and requirements that cannot be fulfilled in many taxa which range from the need of phased haplotypes to recombination maps. The main limitation of these approaches is that many require a set of reference haplotypes—individuals known to be unadmixed representatives of either population—in order to properly infer the origin of each allele in each (non-reference) sample haplotype.

The last family of approaches designed to uncover introgressed loci has focused on the use of relative and absolute levels of divergence measured in genomic windows. Largely such methods have consisted of new summary statistics that capture elements of the expected coalescent genealogy under a model of recent introgression between species. These approaches have the advantage that no donor or recipient populations must be identified a priori. Among the measurements of divergence, *F_ST_* [48] is most commonly used. Another popular point of departure has been the *d_xy_* statistic of Nei and Li [49] which measures the average pairwise distance between alleles sampled from two populations. For instance, Joly et al. [50], Geneva et al. [51] and Rosenzweig et al. [52] use the minimum rather than the mean of these pairwise divergence values, termed *d_min_*. *d_min_* is sensitive to abnormally short branch lengths between alleles drawn from two populations, as would be expected when introgression is recent. Each of these statistics has attractive properties and adequate power in some instances, however no one statistic has perfect sensitivity in every scenario.

Here we introduce a new method for finding introgressed loci based on supervised machine learning that we call FILET (<u>F</u>inding <u>I</u>ntrogressed <u>L</u>oci using <u>E</u>xtra <u>T</u>rees Classifiers). FILET combines a large number of summary statistics (Materials and Methods) that provide complementary information about the shape of the genealogy underlying a region of the genome. These summary statistics include both previously developed statistics (including, but not limited to, those based on *d_min_* and *d_xy_*) as well as 5 new summary statistics that we describe below. Our reasoning for this approach was that by combining many statistics for detecting introgression we should achieve sensitivity to introgression across a larger range of scenarios than accessible to any individual statistic. Buoyed by our recent work showing the power and flexibility of Extra Trees classifiers [53] for population genomic inference [54, 55], we leveraged this machine learning paradigm for identification of introgression. Using simulations we show that FILET is far more powerful and versatile than competing methods for identifying introgressed loci. Further we apply FILET to examine patterns of introgression between *Drosophila simulans* and its island endemic sister taxon *Drosophila sechellia*.

The speciation event that gave rise to the island endemic *Drosophila sechellia* from a *Drosophila simulans*-like ancestor is a textbook example of a specialist species that evolved from a presumably generalist ancestor [56, 57]. Indeed, *D. sechellia* has quite remarkably specialized to breed on the toxic fruit of *Morinda citrifolia* [58], while *D. simulans* (and *D. mauritiana*) do not tolerate the organic volatile compounds in the ripe fruit [59–61]. The genetic and neurological underpinnings of this key ecological difference have been identified, at least in part [62–67] making the *D. simulans*/*D. sechellia* pair one of the most successful cases of genetic dissection of the causes of an ecologically relevant trait. While this is so, the population genetics of divergence between these species has only been examined in the context of either population samples from a handful of loci [68–71] or in the absence of population data [72]. These studies estimated population divergence time between *D. simulans* and *D. sechellia* to be as early as ∼250,000 years ago [72] or as old as ∼413,000 years ago [70]. All population genomic surveys demonstrate that *D. sechellia* harbors little genetic variation in comparison to *D. simulans*, perhaps as a result of a population size crash/founder event from which the population has not recovered [68, 71]. Moreover it has been suggested that what little variation there is in *D. sechellia* shows little population genetic structure among separate island populations in the Seychelles archipelago [71]. Lastly there is some evidence of introgression between each pair of species within the *D. simulans* complex [72], and *D. simulans* and *D. sechellia* have been found to hybridize in the field [73]. Here we characterize the population genetics of divergence between *D. sechellia* and *D. simulans,* combining existing whole-genome sequences from a mainland population of *D. simulans* [74] with newly generated genome sequences from *D. sechellia.* Applying FILET to these data confirms previous reports of introgression between these species and reveals that this gene flow is primarily in the direction of *D. simulans* to *D. sechellia.* Finally, the success of our approach underscores the potential power of supervised machine learning for evolutionary and population genetic inference.

## MATERIALS AND METHODS

### Statistics capturing the population genetic signature of introgression

We set out to assemble a set of statistics that could be used in concert to reliably determine whether a given genomic window had experienced recent gene flow. Several statistics that have been designed to this end ask whether there is a pair of samples exhibiting a lower than expected degree of sequence divergence within the window of interest. The most basic of these is *d_min_*, the minimum pairwise divergence across all cross-population comparisons (S1 Fig; [50]). The reasoning behind *d_min_* is that even if only a single sampled individual contains an introgressed haplotype, *d_min_* should be lower than expected and the introgression event may be detectable. A related statistic is *G_min_*, which is equal to *d_min_*/*d_xy_* [51]; the presence of this term in the denominator is meant to control for variation in the neutral mutation rate across the genome. *RND_min_* accomplishes this by dividing *d_min_* by the average divergence of all sequences from either species to an outgroup sequence [52]. The name of this statistic is derived from its constituent parts, *d_min_*, and *RND* [75].

As described in the following section, we incorporated a number of previously devised statistics into our classification approach, including some of those based on *d_min_*. We also included some novel statistics that we designed to have improved sensitivity to particularly recent introgression. The first of these is defined as:

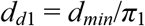

where *π*_1_ is nucleotide diversity [49] in population 1. Similarly, *d_d2_* = *d_min_*/*π*_2_. *d_d_*_1_ and *d_d_*_2_ statistics are so named because they compare *d_min_* to diversity within populations 1 and 2, respectively. The rationale behind these statistics is that, if there has been recent introgression from population 1 into population 2, and at least one sampled chromosome from population 2 contains the introgressed haplotype, then the cross-population pair of individuals yielding the value of *d_min_* should both trace their ancestry to population 1. Thus, the sequence divergence between these two individuals should on average be equal to *π*_1_. Similarly, if there was introgression in the reverse direction *d_min_* would be on the order of *π*_2_. Following similar rationale, we devised two related statistics: *d_d-Rank_*_1_ and *d_d-Rank_*_2_. *d_d-Rank_*_1_ is the percentile ranking of *d_min_* among all pairwise divergences *within* population 1; the value of this statistic should be lower when there has been introgression from population 1 into population 2. *d_d-Rank_*_2_ is the analogous statistic for introgression from population 2 into population 1. We also included a statistic comparing average linkage disequilibrium within populations to average LD within the global population (i.e. lumping all individuals from both species together), as follows:

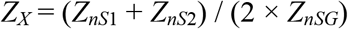

where *Z_nS_*_1_, and Z*_nS_*_2_ measure average LD [76] between all pairs of variants within the window in population 1 and population 2, respectively, and Z*_nSG_* which measures LD within the global population. The reasoning behind this statistic is based on the assumption that, in the presence of gene flow, LD may be elevated within the recipient population(s) but not in the global population. S2 Fig shows that the distributions of these statistics do indeed differ substantially between genealogies with and without introgression (simulation scenarios described below), especially when this introgression occurred recently. In addition to these and other statistics summarizing diversity across the two population samples, we also incorporated several single-population statistics into our classifier (see below), as these may also contain information about recent introgression. For example, separate measures of nucleotide diversity in our two population samples would contain useful information because introgression is expected to increase diversity in the recipient population, especially if the source population was large or if the two populations split long ago.

### Description of FILET classifier

We used a supervised machine learning approach to assign a genomic window to one of three distinct classes on the basis of a “feature vector” consisting of a number of statistics summarizing patterns of variation within the window from two closely related populations. These three classes are: introgression from population 1 into population 2, introgression from population 2 into population 1, and the absence of introgression. Specifically, we used an Extra-Trees classifier [53], which is an extension of random forests [77], an ensemble learning technique that creates a large ensemble of semi-randomly generated binary decision trees [78], before taking a vote among these decision trees in order to decide which class label should be assigned to a given data instance (i.e. genomic window in our case). In an Extra-Trees classifier, the tree building process is even more randomized than in typical random forests: in addition to selecting a random subset of features when generating a tree, the separating threshold for each feature is randomly chosen, rather than selected the threshold that optimally separates the data classes. We require example regions for each class in order to train the Extra-Trees classifier, so we used coalescent simulations to generate these training examples (described below). Our ultimate goal was to detect introgression within 10 kb windows in *Drosophila*, so to train our classifier properly we simulated chromosomal regions approximating this length (simulation details are given below). The target window size could easily be altered by changing the length of the regions simulated for training (i.e. by adjusting the recombination and mutation rates, *θ* and *ρ*).

FILET’s feature vector contains a number of single-population summaries of per-base pair genetic variation: *π*, the variance in pairwise distances among individuals, the density of segregating sites, the density of polymorphisms private to the population, Fay and Wu’s *H* and *θ_H_* statistics [79], and Tajima’s *D* [80]. The feature vector also includes two single-population summary statistics that are not normalized per base pair: *Z_nS_* (which is averaged across all pairs of SNPs), and the number of distinct haplotypes observed in the window. Each feature vector included values of these 9 statistics for each population, yielding 18 single-population statistics in total. In addition, the two-population statistics included in FILET’s feature vector were as follows: *F_ST_* (following ref. [81]), Hudson’s *S*_nn_ [82], per-bp *d_xy_*, per-bp *d_min_*, *G_min_*, *d_d_*_1_, *d_d_*_2_, *d_d-Rank_*_1_, *d_d-Rank_*_2_, *Z_X_*, *IBS_MaxB_*(the length of the maximum stretch of identity by state, or IBS, among all pairwise between-population comparisons), and *IBS_Mean_*_1_ and *IBS_Mean_*_2_ which capture the average IBS tract length when comparing all pairs of sequences within populations 1 and 2, respectively. These IBS statistics are calculated by examining all pairs of individual sequences within a population (or across populations in the case of *IBS_MaxB_*), noting the positions of differences, and examining the distribution of lengths between these positions (as well as between the first position and the beginning of the window and between the last position and the end of the window). Note that we did not include *RND_min_* or other measures such as Patterson’s *D* and *F_4_* statistics [83] that require alignment to one or more additional species along with the focal pair, because we designed FILET so that it would not require outgroup information. We note however that through its use of supervised machine learning, FILET could easily be extended to incorporate such data. Instead, in order to improve robustness to mutational variation, we adopted the approach of drawing the mutation rate from a wide range of values when generating training examples to train FILET to classify data from our *Drosophila* samples (see below). All code necessary to run the FILET classifier (including calculating summary statistics on both simulated and real data sets, training, and classification) along with the full results of our application to *D. simulans* and *D. sechellia* (described below) are available at https://github.com/kern-lab/FILET/.

### Simulated test scenarios

Following Rosenzweig et al. [52], we used the coalescent simulator msmove (https://github.com/geneva/msmove) to simulate data for testing FILET’s power to detect introgression in populations with four different values of *T_D_* (the time since divergence): 0.25×4*N*, 1×4*N*, 4×4*N*, and 16×4*N* generations ago, where *N* is the population size. For each of these simulations the population size was held constant (i.e. the ancestral population size equals that of both daughter populations). We developed a classifier for each of these scenarios of population divergence. Supervised machine learning techniques such as the Extra-Trees classifier require training data consisting of examples from each of the three classes, but in practice a large number of example loci known to have experienced introgression may not be available. We therefore simulated training data sets for each of the four values of *T_D_*. Again following Rosenzweig et al. [52], the relevant parameters for each of these simulations include: *T_M_*, the time since the introgression event, which we drew from {0.01×*T_D_*, 0.05×*T_D_*, 0.1×*T_D_*, 0.15×*T_D_*, …, 0.9×*T_D_*} (i.e. multiples of 0.05×*T_D_* up to 0.9, and also including 0.01×*T_D_*); and *P_M_*, the probability that a given lineage would migrate from the source population to the sink population during the introgression event, which we drew from {0.05, 0.1, 0.15, …, 0.95}. We simulated an equal number of training examples for each combination of these two parameter values for both directions of gene flow, yielding 10^4^ simulations in total for both of these classes, conditioning that each of these instances must have contained at least one migrant lineage. Finally, we simulated an equivalent number of samples without introgression, yielding a balanced training set (10^4^ examples for each class).

Next, we computed feature vectors as described above for each of these training examples, and proceeded with training our Extra-Trees classifiers by conducting a grid search of all training parameters precisely as described in Schrider and Kern [54], and setting the number of trees in the resulting ensemble to 100. All training and classification with the Extra-Trees classifier was performed using the scikit-learn Python library (http://scikit-learn.org; [84]). We also calculated feature importance and rankings thereof by training an Extra-Trees classifier of 500 decision trees on the same training data (using scikit-learn’s defaults for all other learning parameters), and then using this classifier’s “feature_importances_” attribute. Briefly, this feature importance score is the average reduction in Gini impurity contributed by a feature across all trees in the forest, always weighted by the probability of any given data instance reaching the feature’s node as estimated on the training data [85]; this measure thus captures both how well a feature separates data into different classes and how often the feature is given the opportunity to split (i.e. how often it is visited in the forest). The values of these scores are then normalized across all features such that they sum to one.

For each *T_D_*, we evaluated the appropriate classifier against a larger set of 10^4^ simulations generated for each parameter combination along a grid of values of *T_M_* and *P_M_*. The values of *P_M_* were drawn from the same set as those in training as described above, while one additional possible value of *T_M_* was included: 0.001×*T_D_*. Also note that for these simulations we did not require at least one migrant lineage as we had done for training (information that is recorded by msmove). For test simulations with bidirectional migration, we did not require each replicate sample to contain at least one migrant lineage, though we modified msmove to record which samples did in fact experience migration. In addition to test examples for each direction of gene flow, we simulated 10^4^ examples where no migration occurred in order to assess false positive rates. In order to examine the performance of FILET when an unsampled ghost lineage was the source of introgression, we generated test simulations with the same values of *T_D_*, *T_M_*, and *P_M_* as above, but where the source of the introgression was a third, unsampled population that separated from the two sampled populations at time *T_D_*. In all of our simulations, both for training and testing, we set locus-wide population mutation and recombination rates *θ* and *ρ* to 50 and 250, respectively, similar to autosomal values in 10 kb windows in *D. melanogaster* [86] and sampled 15 individuals from each population. We also experimented with different window sizes, simulating training and test data (1,000 replicates for each class for each set) with window sizes corresponding to 1 kb, 10 kb, 5 kb, and 50 kb by multiplying *θ* and *ρ* by the appropriate scalar. When testing the sensitivity of our method on these data, we considered a window to be introgressed if FILET’s posterior probability of the no-introgression class was <0.05, except for the scenario with *T_D_* equal to 16×4*N* generations ago in which case we used a posterior probability cutoff of 0.01, as we found that this step mitigated the elevated false positive rate under this scenario (reducing the rate from >10% to the estimate of 6% shown in S3 Fig). In windows labeled as introgressed, the direction of gene flow was determined by asking which of the two introgression classes had a higher posterior probability. Note that we used the same demographic scenario for both the training and test data for each *T_D_*, and discuss the implications of demographic model misspecification in the Results and Discussion.

In order to compute receiver operator characteristic (ROC) curves we constructed balanced binary training sets composed of 10^4^ examples with no introgression, and 10^4^ examples allowing for introgression (with equal representation to each combination of *T_M_*, *P_M_*, and direction of introgression. The score that we obtained for each test example in order to compute the ROC curve was one minus the posterior probability of no introgression as generated by the Extra-Trees classifier (i.e. the classifier’s estimated probability of introgression, regardless of directionality).

### Comparison to ChromoPainter

We compared FILET’s accuracy to that of ChromoPainter [46], a software program that utilizes a variant of the copying model first proposed by Li and Stephens [87]. In this model each sample haplotype is a mosaic composed of chromosomal segments chosen from a set of possible ancestral haplotypes, allowing for differences caused by mutation and the potential for changes in ancestry at recombination breakpoints. Such an approach can thus be used to predict the ancestry of each individual at each polymorphism—these predictions are referred to as “paintings” by ChromoPainter. To this end we repeated our simulations above but increased the size of the chromosomal segments to 1 Mb by increasing *θ* and *ρ* to 5000 and 25000. In these simulations only gene flow from population 2 to population 1 was allowed, and we modified msmove to record the coordinates of introgressed segments, and to restrict introgression events to those involving segments falling entirely within the central 100 kb of the chromosome. For each combination of *T_M_* and *P_M_* we generated 10 replicate simulations, including 10 replicates without introgression.

We ran ChromoPainter with the following parameters: the “-a 0 0” switch to model each individual haplotype as a mosaic of each other individual rather than using a set of predefined reference haplotypes, “-i 10” and “-ip” options to estimate copying proportions over 10 Expectation-Maximization (EM) iterations, and the default “-s 10” switch to stochastically draw 10 chromosome paintings for each individual from the HMM following EM. We then used the output from ChromoPainter to predict introgressed chromosomal segments as follows:

For each polymorphism, we examine each haplotype *i* among our *n* haplotypes, and record which of the other *n*-1 haplotypes serves as the best ancestor for *i* in each of our 10 paintings. We then examine each individual in population 2 (the recipient population), and count the number of paintings for which the ancestral haplotype is from population 1. If this number is > 5 (i.e. a majority) for any of our individuals in population 2, then we consider this focal polymorphism to be introgressed. If two adjacent polymorphisms are predicted to be introgressed, all sites between them are also considered to be introgressed. If only 1 polymorphism is predicted, then just that one site is considered introgressed. We also produced a more stringent version of these predictions by only retaining introgressed segments consisting of at least 25 consecutive introgressed polymorphisms. Note that ChromoPainter requires base pair positions, and msmove uses an infinite sites model where polymorphisms are located in a continuous space between zero and one. Thus in order to perform this analysis we had to map msmove’s continuous locations to physical locations, which we accomplished by multiplying by 10^6^ and rounding to the nearest available position.

We compared ChromoPainter to a sliding-window application of FILET’s classifier for 10 kb windows (with 1 kb step sizes). We also produced finer-scale FILET predictions using a 1 kb classifier (with 100 bp step sizes) to refine predictions made by the 10 kb classifier: only sliding windows predicted as introgressed by the 1 kb classifier and lying within introgressed segments predicted by the 10 kb classifier were retained as candidates by this version. For the refinement step, FILET’s posterior probability cutoff for introgression was relaxed to 0.5 (i.e. introgression more probable than not); a more lenient cutoff is appropriate here because this classifier was only applied within regions already predicted to be introgressed by the 10 kb classifier.

### Drosophila sechellia collection

*Drosophila sechellia* flies were collected in the islands of Praslin, La Digue, Marianne and Mahé with nets over fresh *Morinda* fruit on the ground. All flies were collected in January of 2012. Flies were aspirated from the nets by mouth (1135A Aspirator – BioQuip; Rancho Domingo, CA) and transferred to empty glass vials with wet paper balls (to provide humidity) where they remained for a period of up to three hours. Flies were then lightly anesthetized using FlyNap (Carolina Biological Supply Company, Burlington, NC) and sorted by sex. Females from the *melanogaster* species subgroup were individualized in plastic vials with instant potato food (Carolina Biologicals, Burlington, NC) supplemented with banana. Propionic acid and a pupation substrate (Kimwipes Delicate Tasks, Irving TX) were added to each vial. Females were allowed to produce progeny and imported using USDA permit P526P-15-02964. The identity of the species was established by looking at the taxonomical traits of the males produced from isofemale lines (genital arches, number of sex combs) and the female mating choice (i.e., whether they chose *D. simulans or D. sechellia* in two-male mating trials).

### Sequence data and variant calling and phasing

We obtained sequence data from 20 *D. simulans* inbred lines [74] from NCBI’s Short Read Archive (BioProject number PRJNA215932). We also sequenced wild-caught outbred *D. sechellia* females (see above) from Praslin (*n*=7 diploid genomes), La Digue (*n*=7), Marianne (*n*=2), and Mahé (*n*=7). These new *D. sechellia* genomes are available on the Short Read Archive (BioProject number PRJNA395473). For each line we then mapped all reads with bwa 0.7.15 using the BWA-MEM algorithm [88] to the March 2012 release of the *D. simulans* assembly produced by Hu et al. [89] and also used the accompanying annotation based on mapped FlyBase release 5.33 gene models [90]. Next, we removed duplicate fragments using Picard (https://github.com/broadinstitute/picard), before using GATK’s HaplotypeCaller (version 3.7; [91–93]) in discovery mode with a minimum Phred-scaled variant call quality threshold (-stand_call_conf) of 30. We then used this set of high-quality variants to perform base quality recalibration (with GATK’s BaseRecalibrator program), before again using the HaplotypeCaller in discovery mode on the recalibrated alignments. For this second iteration of variant calling we used the --emitRefConfidence GVCF option in order to obtain confidence scores for each site in the genome, whether polymorphic or invariant. Finally, we used GATK’s GenotypeGVCFs program to synthesize variant calls and confidences across all individuals and produce genotype calls for each site by setting the --includeNonVariantSites flag, before inferring the most probable haplotypic phase using SHAPEIT v2.r837 [94]. The genotyping and phasing steps were performed separately for our *D. simulans* and *D. sechellia* data, and for each of step in the pipeline outlined above we used default parameters unless otherwise noted. In order to remove potentially erroneous genotypes (at either polymorphic or invariant sites), we considered genotypes as missing data if they had a quality score lower than 20, or were heterozygous in *D. simulans*. After throwing out low-confidence genotypes, we masked all sites in the genome missing genotypes for more than 10% of individuals in either species’ population sample, as well as those lying within repetitive elements as predicted by RepeatMasker (http://www.repeatmasker.org). Only SNP calls were included in our downstream analyses (i.e. indels of any size were ignored). These phased and masked data are available at https://zenodo.org/record/1166021.

### Demographic inference

Having obtained genotype data for our two population samples, we used *∂*a*∂*i [95] to model their shared demographic history on the basis of the folded joint site frequency spectrum (downsampled to *n*=18 and *n*=12 in *D. simulans* and *D. sechellia*, respectively); using the folded spectrum allowed us to circumvent the step of producing whole genome alignments to outgroup species in *D. simulans* coordinate space in order to attempt to infer ancestral states. We used an isolation-with-migration (IM) model that allowed for continual exponential population size change in each daughter population following the split. This model includes parameters for the ancestral population size (*N_anc_*), the initial and final population sizes for *D. simulans* (*N_sim_0_* and *N_sim_*, respectively), the initial and final sizes for *D. sechellia* (*N_sech_0_* and *N_sech_*, respectively), the time of the population split (*T_D_*), the rate of migration from *D. simulans* to *D. sechellia* (*m_sim→sech_*), and the rate of migration from *D. sechellia* to *D. simulans* (*m_sech→sim_*). We also fit our data to a pure isolation model that was identical to our IM model but with *m_sim→sech_* and *m_sech→sim_* fixed at zero. Finally, we fit our data to an admixture model identical to the isolation model but with the addition of two parameters: the time of a pulse admixture event from *D. simulans* into *D. sechellia* (*T_AD_*) and the proportion of individuals in *D. sechellia* migrating from *D. simulans* during this event (*P_AD_*). Our optimization procedure consisted of an initial optimization step using the Augmented Lagrangian Particle Swarm Optimizer [96], followed by a second step of optimization refining the initial model using the Sequential Least Squares Programming algorithm [97], both of which are included in the pyOpt package for optimization in Python (version 1.2.0; [98]) as in Schrider et al. [99]. We performed ten optimization runs fitting both of these models to our data, each starting from a random initial parameterization, and assessed the fit of each optimization run using the AIC score. Code for performing these optimizations can be obtained from https://github.com/kern-lab/miscDadiScripts, wherein 2popIM.py, 2popIsolation.py, 2popIsolation_admixture.py fit the IM, isolation, and admixture models described above, respectively. For scaling times by years rather than numbers of generations, we assumed a generation time of 15 gen/year as has been estimated in *D. melanogaster* [100].

### Training FILET to detect introgression between *D. simulans* and *D. sechellia*

Having obtained a demographic model that provided an adequate fit to our data, we set out to simulate training examples under this demographic history for each of our three classes (i.e. no migration, migration from *D. simulans* to *D. sechellia*, and from *D. sechellia* to *D. simulans*). For training examples including introgression, *T_M_* was drawn uniformly from the range between zero generations ago and *T_D_*/4, while *P_M_* raged uniformly from (0, 1.0]. In addition, in order to make our classifier robust to uncertainty in other parameters in our model, for each training example we drew values of each of the remaining parameters from [*x*−(*x*/2), *x*+(*x*/2)], where *x* is our point estimate of the parameter from *∂*a*∂*i. In addition to the parameters from our demographic model (*T_D_*, *ρ*, *N_anc_*, *N_sim_*, and *N_sech_*), these include the population mutation rate *θ*=4*Nµ* (where *µ* was set to 3.5×10^-9^), and the ratio of *θ* to the population recombination rate *ρ* (which we set to 0.2). Continuous migration rates were set to zero (i.e. the only migration events that occurred were those governed by the *T_M_* and *P_M_* parameters, and the no-migration examples were truly free of migrants). In total, this training set comprised of 10^4^ examples from each of our three classes.

As described above, we masked genomic positions having too many low confidence genotypes or lying within repetitive elements (described above) before proceeding with our classification pipeline. While doing so, we recorded which sites were masked within each 10 kb window in the genome that we would later attempt to classify. In order to ensure that our masking procedure affected our simulated training data in the same manner as our real data, for each simulated 10 kb window we randomly selected a corresponding window from our real dataset and masked the same sites in the simulated window that had been masked in the real one. We then trained our classifier in the same manner as described above.

In order to ensure that this classifier would indeed be able to reliably uncover loci experiencing recent gene flow between our two populations, we assessed its performance on simulated test data. First, we applied the classifier to test examples simulated under this same model (again, 10^4^ for each class). Next, to address the effect of demographic model misspecification, we applied our classifier to an isolation model with a different parameterization and no continuous size change in the daughter populations. For this model we simply set *N_sim_* and *N_sech_* to *π_sim_*/4*µ* and *π_sech_*/4*µ*, respectively, where *π* for a species is the average nucleotide diversity among all windows included in our analysis after filtering, and *µ* was again set to 3.5×10^-9^. We then set *N_anc_* to be equal to *N_sim_*, and set *T* to *d_xy_*/(2*µ*) − 2*N_anc_* generations where *d_xy_* is the average divergence between *D. simulans* and *D. sechellia* sequences across all windows. This latter value is simply the expected TMRCA for cross-species pairs of genomes, minus the expected waiting time until coalescence during the one-population (i.e. ancestral) phase of the model. This simple model thus produces samples with similar levels of nucleotide diversity for the two daughter populations as those produced under our IM model, but that would differ in other respects (e.g. the joint site frequency spectrum and linkage disequilibrium, which would be affected by continuous population size change after the split). For both test sets we masked sites in the same manner as for our training data before running FILET.

### Classifying genomic windows with FILET

We examined 10 kb windows in the *D. simulans* and *D. sechellia* genomes, summarizing diversity in the joint sample with the same feature vector as used for classification (see above), ignoring sites that were masked as described above. We omitted from this analysis any window for which >25% of sites were masked, and then applied our classifier to each remaining window, calculating posterior class membership probabilities for each class. We then used a simple clustering algorithm to combine adjacent windows showing evidence of introgression into contiguous introgressed elements: we obtained all stretches of consecutive windows with >90% probability of introgression as predicted by FILET (i.e. the probability of no-introgression class <10%), and retained as putatively introgressed regions those that contained at least one window with >95% probability of introgression. In order to test for enrichment of these introgressed regions for genic/intergenic sequence or particular Gene Ontology (GO) terms from the FlyBase 5.33 annotation release [90], we performed a permutation test in which we randomly assigned a new location for each cluster or introgressed windows (ensuring the entire permuted cluster landed within accessible windows of the genome according to our data filtering criteria). We generated 10,000 of these permutations.

## RESULTS AND DISCUSSION

### FILET detects introgressed loci with high sensitivity and specificity

We sought to devise a bioinformatic approach capable of leveraging population genomic data from two related population samples to uncover introgressed loci with high sensitivity and specificity. In the Materials and Methods, we describe several previous and novel statistics designed to this end. However, rather than preoccupying ourselves with the search for the ideal statistic for this task, we took the alternative approach of assembling a classifier leveraging many statistics that would in concert have greater power to discriminate between introgressed and non-introgressed loci. Supervised machine learning methods have proved highly effective at making inferences in high-dimensional datasets and are beginning to make inroads in population genetics [101]. In this vein, we designed FILET, which uses an extension of random forests called an Extra-Trees classifier [53]. We previously succeeded in applying Extra-Trees classifiers for a separate population genetic task—finding recent positive selection and discriminating between hard and soft sweeps [54, 55]—though other methods such as support vector machines [102] or deep learning [103] could also be applied to this task.

Briefly, FILET assigns a given genomic window to one of three distinct classes—recent introgression from population 1 into population 2, introgression from population 2 into 1, or the absence of introgression—on the basis of a vector of summary statistics that contain information about the two-population sample’s history. This feature vector contains a variety of statistics summarizing patterns of diversity within each population sample, as well as a number of statistics examining cross-population variation (see Materials and Methods for a full description). FILET must be trained to distinguish among these three classes, which we accomplish by supplying 10,000 simulated example genomic windows of each class, with each example represented by its feature vector. Because we expect that the majority of introgression events to be non-adaptive, these simulations did not include natural selection. Once this training is complete, FILET can then be used to infer the class membership of additional genomic windows, whether from simulated or real data.

We began by assessing FILET’s power on a number of simulated datasets, examining windows roughly equivalent to 10 kb in length in *Drosophila* (Materials and Methods). In particular, because the power to detect introgression depends on the time since their divergence, *T_D_*, we measured FILET’s performance under four different values of *T_D_*, training a separate classifier for each. In Figure 1 (*T_D_*=0.25×4*N*) and S3 Fig (*T_D_* values of 1, 4, and 16×4*N*), we compare FILET’s power to that of two related statistics that have been devised to detect introgressed windows, *d_min_* and *G_min_* (Materials and Methods). These figures show that FILET has high sensitivity to introgression across a much wider range of introgression timings (*T_M_*) and intensities (*P_M_*) than either of these statistics under each value of *T_D_*, and that this disparity is amplified dramatically for smaller values of *T_D_*. Furthermore, these figures demonstrate that FILET infers the correct directionality of recent introgression with high accuracy, whereas *d_min_* and *G_min_* contain no information about the direction of gene flow. Finally, FILET does not appear especially sensitive when the source of gene flow is an unsampled ghost population rather than one of the two sequenced populations (S4 Fig), though it could potentially be trained to detect such cases if desired.

**Fig. 1.**
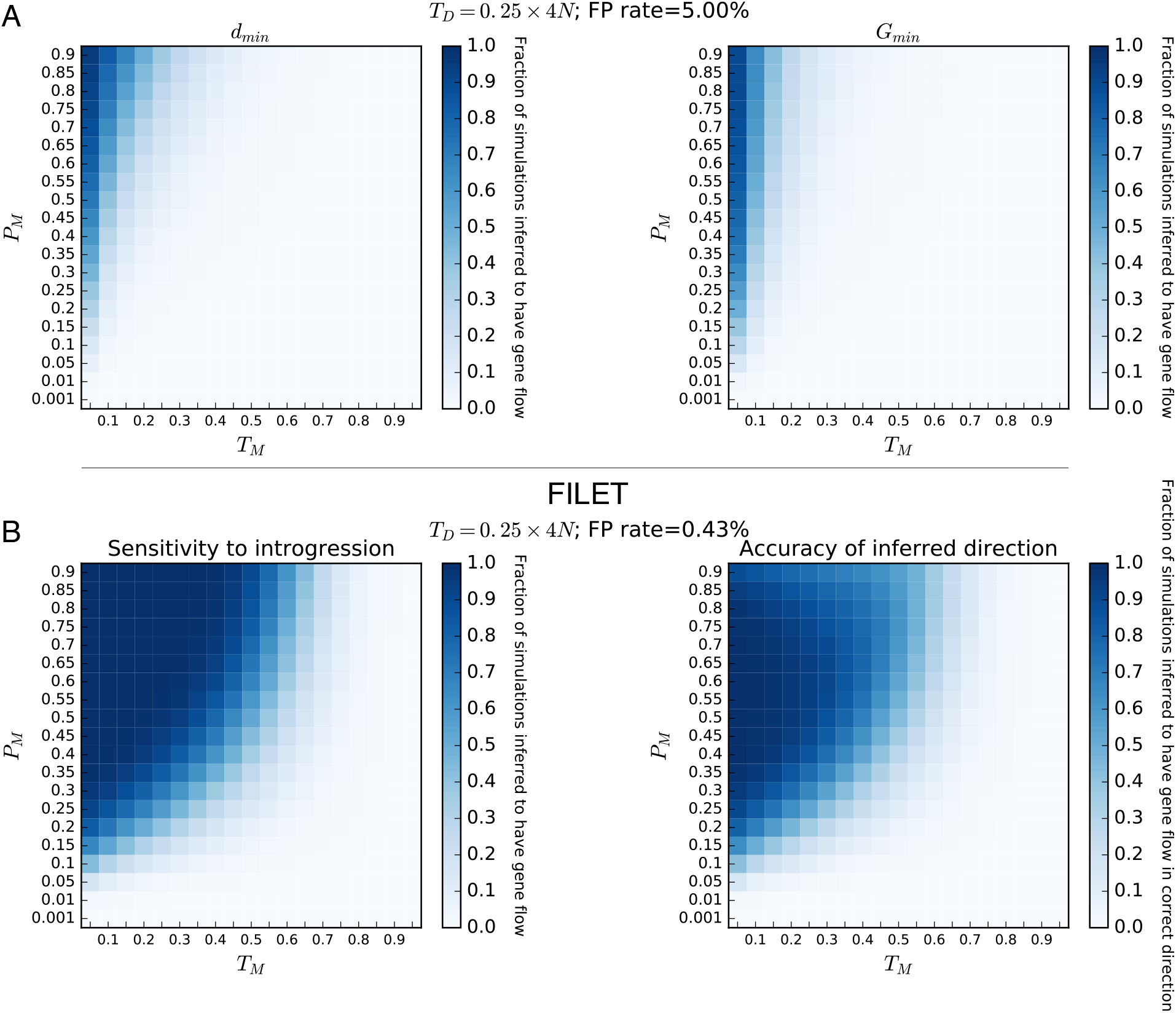
Heatmaps showing several methods’ sensitivity to detect introgression. We show the fraction of simulated genomic regions with introgression occurring under various combinations of migration times (*T_M_*, shown as a fraction of the population divergence time *T_D_*) and intensities (*P_M_*, the probability that a given lineage will be included in the introgression event) that are detected successfully by each method. (*A*) Accuracy of *d_min_* and *G_min_* statistics, where a simulated region is classified as introgressed if the values of these statistics are found in the lower 5% tail of the distribution under complete isolation (from simulations). Thus, the false positive rate is fixed at 5%. (*B*) The accuracy of FILET on these same simulations. On the left we show the fraction of regions correctly classified as introgressed (compare to panel *A*). On the right, we show the fraction of all simulated regions that are not only classified as introgressed, but also for which the direction of gene flow was correctly inferred (i.e. if the direction is inferred with 100% accuracy for a given cell in the heatmap, the color shade of that cell will be identical to that in the heatmap on the left). The false positive rate, as determined from applying FILET to a simulated test set with no migration, is also shown.

We also note that for *d_min_* and *G_min_* we established 95% significance thresholds from our simulated training data without introgression, thereby achieving a false positive rate of 5%. In order to assess FILET’s false positive rate, we classified a set of test simulations without introgression and found that FILET’s false positive rate was considerably lower (Figure 1 and S3 Fig) except for our largest value of *T_D_*, where it was initially higher (0.4% for *T_D_*=0.25×4*N* but >10% for *T_D_*=16×4*N*), despite our selection of a posterior probability cutoff of 95% (Methods). This illustrates an important issue with posterior probability estimates produced by supervised machine learning methods: they may occasionally be miscalibrated. We therefore adjusted the cutoff for the *T_D_*=16×4*N* scenario (to 99% probability of introgression) which lowered our false positive rate to 6% as shown in S3 Fig. Thus, when an appropriate posterior probability cutoff is chosen—a task that can be performed in a straightforward manner by testing on simulated data— FILET achieves much greater sensitivity to introgression than *d_min_* and *G_min_* often at a much lower false positive rate. We also demonstrate the FILET’s greater power than these statistics via ROC curves (S5 Fig), where it outperforms each statistic under each scenario. Specifically, the difference in power between FILET and *d_min_* is dramatic for smaller values of *T_D_* (area under curve, or AUC, of 0.85 versus 0.73 when *T_D_*=0.25×4*N* for FILET and *d_min_*, respectively) but comparatively miniscule for our largest *T_D_* (AUC of 0.94 versus 0.93 when *T_D_*=16×4*N*). It therefore appears that FILET’s performance gain relative to single statistics is highest for the more difficult task of finding introgression between very recently diverged populations, while for the easier case of detecting introgression between highly diverged populations some single statistics may perform nearly as well.

We also experimented with smaller training sets, finding similar classification power (measured by AUC) as above when we trained FILET using only 1000 or even 100 simulated examples per class (S6 Fig), though in the latter case estimated class posterior probabilities were far less accurate. In addition, we examined the effect of altering the target window size used when training and testing FILET (S7 Fig).

### A comparison of the power and resolution of FILET and ChromoPainter

Methods designed to uncover changes in ancestry along a recombining chromosome within admixed populations can also be used to recover introgressed regions. To this end we used ChromoPainter [46] which has the advantage of not requiring a set of “reference haplotypes” known to be free of introgression from each population, and can instead predict for each haplotype, which of all the other haplotypes in the sample (from either population) is most closely related. We simulated two-population samples for 1 Mb chromosomes where introgression from population 2 to population 1 was allowed in the central 100 kb window, and used ChromoPainter to identify introgressed loci (see Methods). We then ran FILET on these simulations, this time using a sliding-window approach to detect introgressed segments (Methods).

Figure 2 shows that FILET has substantially higher sensitivity than ChromoPainter— summing across the entire parameter space (including many scenarios where introgression is quite difficult to detect) FILET recovered 27.7% of introgressed base pairs compared to 19.4% for ChromoPainter—while having a roughly 20-fold lower false positive rate (0.42% for FILET versus 9.31% for ChromoPainter). For scenarios with more ancient and less intense introgression, we did observe somewhat higher sensitivity in ChromoPainter’s predictions. However, this seems to be driven largely by ChromoPainter’s propensity to identify a larger fraction of base pairs as introgressed regardless of their true ancestry, as evidenced by its higher false positive rate. To demonstrate this further we show for the positive predictive value (the number of base pairs correctly predicted to be introgressed divided by the total number of base pairs predicted to be introgressed) for each method in S8 Fig. This figure shows that FILET’s positive predictive is consistently far higher than ChromoPainter’s. We sought to improve this by adopting a more stringent threshold for ChromoPainter’s predictions, requiring at least 25 adjacent polymorphisms to be called introgressed in order to retain the candidate region. This approach did succeed at reducing the false positive rate to 1.15%, though this is still substantially higher than FILET’s, but this improvement came at the cost of ChromoPainter’s sensitivity, which was reduced to 8.6%, roughly one-third that of FILET (Figure 2 and S8 Fig). We also tried an intermediate threshold (5 polymorphisms), but did not observe a substantial increase in specificity compared to our initial more lenient approach (8.49% false positive rate). Thus, while we cannot rule out that it may be possible to devise a method to leverage ChromoPainter’s model to predict introgression that exceeds the performance of our application of ChromoPainter here, our results suggest that it is unlikely that such an approach would eclipse the performance of FILET. We note that ChromoPainter does have the advantage of not requiring simulated training data. ChromoPainter also has the potential to identify donor and recipient haplotypes, which FILET currently does not, but the far greater power of FILET demonstrated above will make it preferable to many researchers who are interested in identifying introgressed regions. Moreover our above results imply that predictions of the span and origin of introgressed haplotypes made directly from ChromoPainter’s output may not always be particularly reliable.

**Fig. 2.**
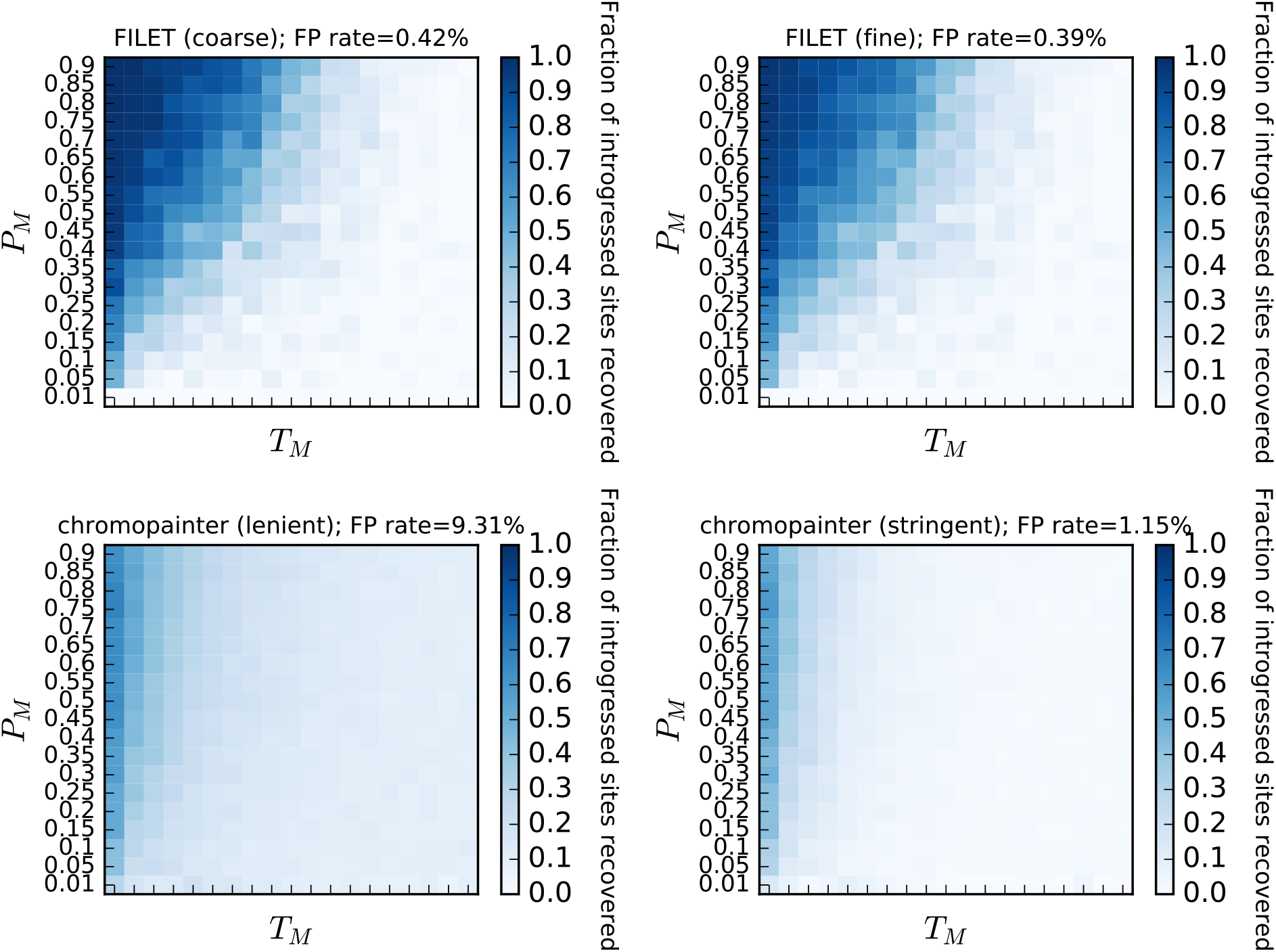
A comparison of the power and resolution of FILET and ChromoPainter using simulations of a 1 Mb chromosome where introgression was allowed within the central 100 kb region. As in Figure 1, the population split time was set to *N* generations ago, and the darkness of the heatmap shows sensitivity to introgression. Unlike Figure 1, here we are measuring sensitivity at the level of the individual base pair rather than evaluating the question of whether a window at large was recovered as containing introgressed alleles. The “coarse” version of FILET refers to a FILET classifier trained to detect introgression in 10 kb windows, which was applied to sliding windows (1 kb step size) across the chromosome. The “fine” version of FILET applied a classifier trained on 1 kb windows to sliding windows (100 bp step size) within those regions classified as introgressed by the FILET classifier. The lenient version of ChromoPainter required evidence of introgression at a single SNP to identify introgression, while the stringent version required candidate regions to contain at least 25 consecutive SNPs supporting introgression.

It is important to note that in the above simulations many introgressed regions will be considerably smaller than our 10 kb window size. This fact, combined with our use of accuracy measurements counting the number of individual base pairs correctly classified, makes the results presented above useful for evaluating FILET’s resolution and the impact of window size on our predictions. By these measures FILET outperforms ChromoPainter, which does not use windows and is only limited in scale by the density of polymorphisms. This suggests that when using sliding windows FILET is able to achieve adequate resolution regardless of its use of a predefined window size. Nonetheless we sought to improve our resolution further by using a finer-scale FILET classifier trained on 1 kb windows as described above to refine the location of putatively introgressed regions identified by the 10 kb classifier (see Methods). While this did marginally reduce our false positive rates and increase our positive predictive values (see Figure 2 and S8 Fig), sensitivity was also somewhat reduced (to 25.7%; Figure 2). The relatively minor effect of adding this refinement step reinforces the notion that a predefined window size is not a major hindrance to our methods’ effectiveness. Thus for most applications a window size that yields enough polymorphisms to reliably calculate the statistics included in our feature vectors may suffice.

Overall, FILET detects introgressed regions with greater power and resolution than ChromoPainter, a method designed to detect ancestry tracks along recombining chromosomes. However we note that many methods of this class exist, and it is possible that some may achieve greater accuracy in some circumstances (e.g. if reference haplotype panels are used).

### Sensitivity to continuous gene flow

While FILET is designed for identifying particular genomic windows that experienced introgression as a result of a pulse migration event, genomic windows with genealogies that include introgression may of course also be produced by continuous migration, with the timing of geneflow varying from window to window. We therefore simulated genomic windows experiencing a variety of bidirectional migration rates under each of our values of *T_D_* and recorded the fraction of windows for which our sampled individuals contained at least one migrant lineage. Next, for each simulated window we applied the FILET classifier trained under the appropriate divergence time as described above, recording the fraction of windows with at least one migrant lineage that were classified as introgressed by FILET. The results of these tests (S1 Table) show that for each value of *T_D_*, the lowest bidirectional migration rates that we tested do not produce migrant lineages, while higher rates will produce a small to modest fraction migrants, most of which are undetected (e.g. when *m*=0.01, 23% and 59% of windows contain at least one migrant when *T_D_*=0.25 and *T_D_*=1, respectively, but <5% are detected by FILET). Thus, FILET, as currently trained, may not be sensitive to gene flow produced by low levels of continuous migration. However, as the migration rate increases further, more and more of these migrant windows will be detected (e.g. when *m*=1, 70% and 100% of windows are detected as migrants by FILET when *T_D_*=0.25 and *T_D_*=1, respectively).

### Ranking the importance of statistics for detecting introgression

Although our goal was to use a set of statistics to perform more accurate inference than possible using individual ones, another benefit of our Extra-Trees approach is that it also allows for a natural way to evaluate the extent to which different statistics are informative under different scenarios of introgression. To this end, we used the Extra-Trees classifier to calculate feature importance, which captures the ability of each statistic to separate the data into its respective classes (Materials and Methods). We find that for our lowest *T_D_* (a split *N* generations ago) the top four features, all with similar importance, are the density of private alleles in population 1, the density of private alleles in population 2, *d_d-Rank_*_1_, and *d_d-Rank_*_2_. For our next-lowest *T_D_*(4*N* generations ago), the top four, again with similar importance score estimates, are *F_ST_*, *Z_X_*, *d_d_*_1_, and *d_d_*_2_. Thus our newly devised *d_d_* and *Z_X_* statistics seem to be particularly informative in the case of recent introgression between closely related populations. For the larger values of *T_D_*, *d_xy_* and *d_min_* rise to prominence. The complete lists of feature importance for each *T_D_* are shown in S2 Table.

The exceptional accuracy with which FILET uncovers introgressed loci underscores the potential for machine learning methods to yield more powerful population genetic inferences than can be achieved via more conventional tools which are often based on a single statistic. Indeed, machine learning tools have been successfully leveraged in efforts to detect recent positive selection [54, 104–107], to infer demographic histories [108], or even to perform both of these tasks concurrently [109].

### Joint demographic history of *D. simulans* and *D. sechellia*

As described in the Materials and Methods, we used publically available *D. simulans* sequence data [74], and collected and sequenced a set of *D. sechellia* genomes. We mapped reads from these genomes to the *D. simulans* assembly [89], obtaining high coverage >28× for each sequence (see sampling locations, mapping statistics, and Short Read Archive identifier information listed in S3 Table). We do not expect that our reliance on the *D. simulans* assembly resulted in any appreciable bias, as reads from our *D. sechellia* genomes were successfully mapped to the reference genome at nearly the same rate as reads from *D. simulans* (S3 Table).

After completing variant calling and phasing (Materials and Methods), we performed principal components analysis on intergenic SNPs at least 5 kb away from the nearest gene in order to mitigate the bias introduced by linked selection [99, 110, 111]. While this is unlikely to completely eliminate the confounding effect of linked selection in *Drosophila*, the fraction of mutations that are deleterious is far greater in coding regions than in intergenic regions (∼90% versus <50% according to [112]); thus it is reasonable to presume that the impact of linked selection is reduced several kilobases away from genes [113]. We observed evidence of population structure within *D. sechellia*. In particular, the samples obtained from Praslin clustered together, while all remaining samples formed a separate cluster (S9 FigA). Running fastStructure [114] on this same set of SNPs yielded similar results: when the number of subpopulations, *K*, was set to 2 (the optimal value for *K* selected by fastStructure’s chooseK.py script), our data were again subdivided into Praslin and non-Praslin clusters. Indeed, across all values of *K* between 2 and 8, fastStructure’s results were suggestive of marked subdivision between Praslin and non-Praslin samples, and comparatively little subdivision within the non-Praslin data (S9 FigB). This surprising result differs qualitatively from previous observations from smaller numbers of loci [71, 115], and underscores the importance of using data from many loci—preferably intergenic and genome-wide—in order to infer the presence or absence of population structure.

Next, we examined the site frequency spectra of the Praslin and non-Praslin clusters, noting that both had an excess of intermediate frequency alleles in comparison to that of the *D. simulans* dataset (S10 Fig), in line with our expectations of *D. sechellia’s* demographic history. We also note that the Praslin samples contained far more variation (50,243 sites were polymorphic within Praslin) than non-Praslin samples (4,108 SNPs within these samples). This difference in levels of variation may reflect a much lesser degree of population structure and/or inbreeding on the island of Praslin than across the other islands, or may result from other demographic processes. Additional samples from across the Seychelles would be required to address this question. In any case, in light of this observation we limited our downstream analyses of *D. sechellia* sequences to those from Praslin.

Because we required a model from which to simulate training data for FILET, we next inferred a joint demographic history of our population samples using *∂*a*∂*i [95]. In particular, we fit three demographic models to our dataset: an isolation-with-migration (IM) model allowing for continuous population size change and migration following the population divergence, an isolation model with the same parameters but fixing migration rates at zero, and an isolation model with one burst of pulse migration from *D. simulans* into *D. sechellia* (Materials and Methods). In S4 Table we show our model optimization results, which show clear support for the IM model over the other models. The IM model that provided the best fit to our data (Figure 3A) includes a much larger population size in *D simulans* than *D. sechellia* (a final size of 9.3×10^6^ for *D. simulans* versus 2.6×10^4^ for *D. sechellia*), consistent with the much greater diversity levels in *D. simulans* [10, 71]. This model also exhibits a modest rate of migration, with a substantially higher rate of gene flow from *D. simulans* to *D. sechellia* (2×*N_anc_m*=0.086) than vice-versa (2×*N_anc_m*=0.013). Thus, the results of our demographic modeling are consistent with the observation of hybrid males in the Seychelles [73], and the possibility of recent introgression between these two species across a substantial fraction of the genome (see refs. [72, 116]).

**Fig. 3.**
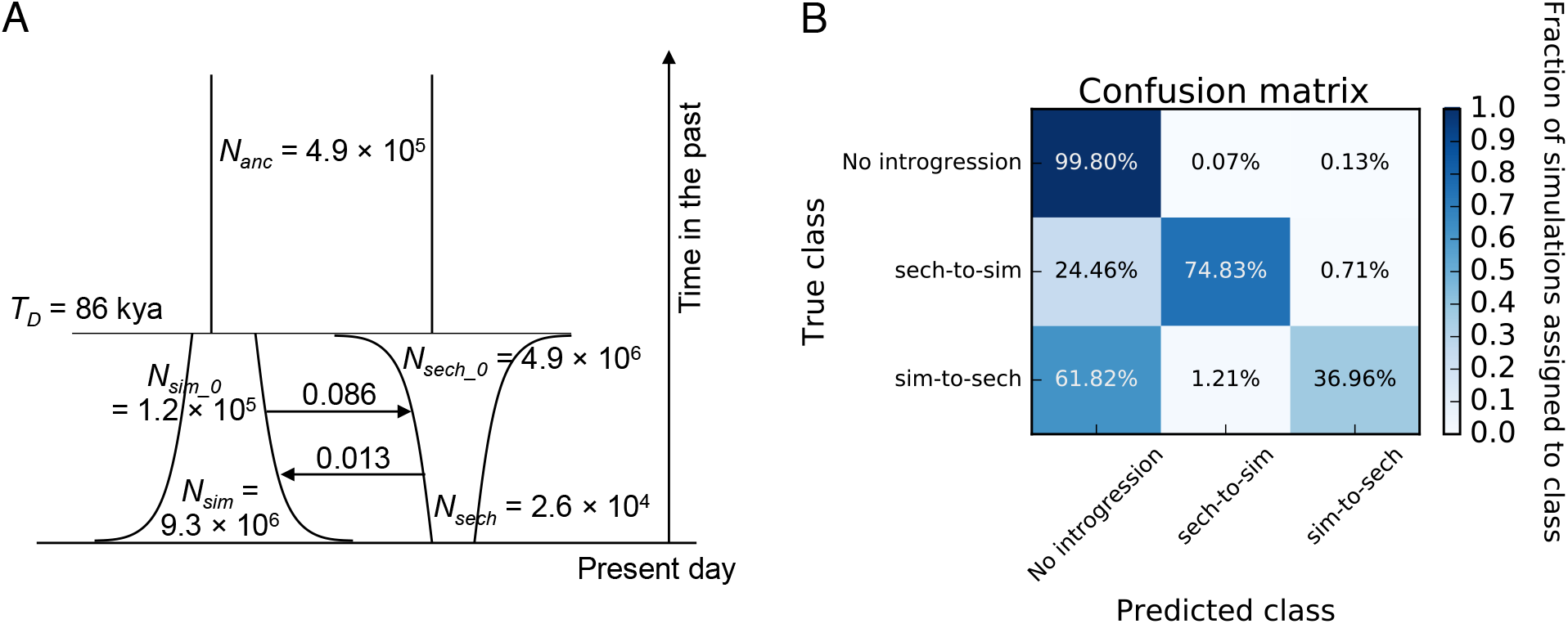
Inferred joint population history of *D. simulans* and *D. sechellia*, and power to detect introgression under this model. (*A*) The parameterization of our best-fitting demographic model. Migration rates are shown by arrows, and are in units of 2×*N_anc_m*, where *m* is the probability of migration per individual in the source population per generation. (*B*) Confusion matrix showing FILET’s classification accuracy under this model as assessed on an independent simulated test set. Perfect accuracy would be 100% along the entire diagonal from top-left to bottom-right, and the false positive rate is the sum of top-middle and top-right cells.

An interesting characteristic of the model shown in Figure 3A is that, assuming 15 generations per year, the estimated time of the *D. simulans*-*D. sechellia* population split is ∼86 kya, or 1.3×10^6^ generations ago. This contrasts with a recent estimate of 2.5×10^6^ generations ago from Garrigan et al. [72] which was based on single genomes rather than population genomic data, but did account for ancestral polymorphism, as did estimates from Obbard et al. [117] which yielded even older split times. Supporting our inference, we note that our average intergenic cross-species divergence of 0.017 yields an average TMRCA of ∼2.5×10^6^ generations ago, assuming a mutation rate of 3.5×10^-9^ mutations per generation as observed in *D. melanogaster* [112, 118], and this estimate would include the time before coalescence in the ancestral population. Unless the mutation rate the *D. simulans* species complex is substantially lower than in *D. melanogaster*, a population split time of 2.5×10^6^ generations ago therefore seems unlikely given that the ancestral population size (and therefore the period of time between the population divergence and average TMRCA) was probably large (>500,000 by our estimate). Thus, we conclude that the *D. simulans* and *D. sechellia* populations may have diverged more recently than previously appreciated, perhaps within the last 100,000 years.

Although the specific parameterization of our model should be regarded as a preliminary view of these species’ demographic history that is adequate for the purposes of training FILET, future efforts with larger sample sizes will be required to refine this model. That being said, the basic features of this model—a much larger *D. simulans* population size than *sechellia*, and a fairly large ancestral population size—are unlikely to change qualitatively.

### Widespread introgression from *D. simulans* to *D. sechellia*

#### Accuracy and robustness of FILET under estimated model

Having obtained a suitable model of the *D. simulans*-*D. sechellia* joint demographic history, we proceeded to simulate training data and train FILET for application to our dataset (Materials and Methods). After training FILET and applying it to simulated data under the estimated demographic model, we find that we have good sensitivity to introgression (56% of windows with introgression are detected, on average), and a false positive rate of only 0.2% (Figure 3B). Thus, while we may miss some introgressed loci, we can have a great deal of confidence in the events that we do recover. Our feature rankings for this classifier are included in S2 Table—under this scenario the most informative feature is our newly devised *d_d-sim_*. Note that we achieve high accuracy despite some of the difficulties presented by the demographic model in Figure 3A, most notably the asymmetry in effective population sizes between our two species. Indeed, because our method is trained under this demographic history, such characteristics of genealogies produced under the assumed demographic history (such as asymmetry in *π*) with and without introgression become the signals used by FILET to make its classifications.

As shown in Figure 3B we find that this classifier has greater sensitivity to introgression from *D. sechellia* to *D. simulans* than vice-versa. The cause of a stronger signal of *D. sechellia*→*D. simulans* introgression can be understood from a consideration of the *d_min_* statistic under each of our three classes. When there is no introgression, *d_min_* will be similar to the expected divergence between *D. simulans* and *D. sechellia*; when there is introgression from *D. simulans* to *D. sechellia*, we may expect *d_min_* to be proportional to *π_sim_*, which may only be a moderate reduction relative to the no-introgression case given the large population size in *D. simulans*; when there is introgression from *D. sechellia* to *D. simulans* then *d_min_* is proportional to *π_sech_* which is dramatically lower than the expectation without introgression. While many of our statistics do not rely on *d_min_*, this example illustrates an important property of the genealogies which include introgression from *D. sechellia* to *D. simulans* that would make them easier to detect than gene flow in the reverse direction.

We also tested this classifier’s performance on a different demographic scenario (S4 Table) in order to examine the effect of model misspecification during training. In particular, we devised a simple island model with two population sizes: a larger size for *D. simulans* and the ancestral population (7.6×10^5^), and a smaller size for *D. sechellia* (5.7×10^4^) with a split time of ∼59 kya. Our simple procedure for estimating these values is described in the Materials and Methods. Again, we find that we have good power to detect introgression with a very low false positive rate (0.28%; S11 Fig). Although there are myriad incorrect models that we could test FILET against, this example suggests that FILET’s performance is robust to at least some scenarios of demographic misspecification.

#### Application to population genomic data

We applied FILET to 10,185 non-overlapping 10 kb windows that passed our data quality filters (101.85 Mb in total, or 86.7% of the five major chromosome arms; Materials and Methods). FILET classified 267 windows as introgressed with high-confidence, which we clustered into 94 contiguous regions accounting for 2.93% of the accessible portion of the genome (2.99 Mb in total; Materials and Methods). This finding is qualitatively similar to a previous estimate (4.6%) by Garrigan et al. [72] based on comparisons of single genomes from each species in the *D. simulans* complex. Unlike this previous effort, FILET is able to infer the directionality of introgression with high confidence (Figure 3B), and we find evidence that the majority of this introgression has been in the direction of *D. simulans* to *D. sechellia*: only 21 of the 267 (7.9%) putatively introgressed windows were classified as introgressed from *sechellia* to *D. simulans*. This finding is not a result of a detection bias, as we have greater power to detect gene flow from *D. sechellia* to *D. simulans* than in the reverse direction. Given that our *D. simulans* sequences are from the mainland, one interpretation of this result is that although there has been recent gene flow from *D. simulans* into the Seychelles, where *D. simulans* and *D. sechellia* occasionally hybridize, there does not appear to be an appreciable rate of back-migration to the mainland of *D. simulans* individuals harboring haplotypes donated from *D. sechellia*. On the other hand, *D. sechellia* alleles may often be purged from *D. simulans* by natural selection. This may be in part due to the reduced ecological niche size of *D. sechellia*, such that any alleles which may introgress into *D. simulans* and lead to preference for or resistance to *Morinda* fruit may prove deleterious in other environments. More generally, *D. sechellia* haplotypes introgressing into *D. simulans* may harbor more deleterious alleles due to their smaller population size, which will be more effectively purged in the larger *D. simulans* population if mutations are not fully recessive [28]. Tests of these hypotheses will have to wait for a population sample of genomes from *D. simulans* collected in the Seychelles.

We asked whether our candidate introgressed loci were enriched for particular GO terms using a permutation test (Materials and Methods), finding no such enrichment. We did observe a deficit in the number of genes either partially overlapping or contained entirely within introgressed regions in our true set versus the permuted set; although a paucity of introgressed genes would be consistent with introgressed functional sequence often being deleterious, this difference was not significant (297 vs. 373.2, respectively; *P*=0.083; one-sided permutation test).

One notable introgressed region on 3R that FILET identified had been previously found by Garrigan et al. as containing a 15 kb region of introgression. We show that gene flow in this region actually extends for over 200 kb (Figure 4). When Brand et al. [119] sequenced the 15 kb region originally flagged by Garrigan et al. in a number of *D. simulans* and *D. sechellia* individuals, they also uncovered evidence of a selective sweep in *D. sechellia* originating from an adaptive introgression from *D. simulans*. Our data set also supports the presence of an adaptive introgression event at this locus: a 10 kb window lying within the putative sweep region (highlighted in Figure 4) is in the lower 5% tail of both *d_min_* (consistent with introgression) and *π_sech_*(consistent with a sweep in *sechellia*); this is the only window in the genome that is in the lower 5% tail for both of these statistics. This region contains two ionotropic glutamate receptors, *CG3822* and *Ir93a*, which may be involved in chemosensing among other functions [120], and the latter of which appears to play a role in resistance to entomopathogenic fungi [121]. Also near the trough of variation within *D. sechellia* are several members of the *Turandot* gene family, which are involved in humoral stress responses to various stressors including heat, UV light, and bacterial infection [122, 123], and perhaps parasitoid attack as well [124]. On the other hand, Brand et al. [119] hypothesize that the target of selection may be a transcription factor binding hotspot between *RpS30* and *CG15696*, and the phenotypic target of this sweep remains unclear.

**Fig. 4.**
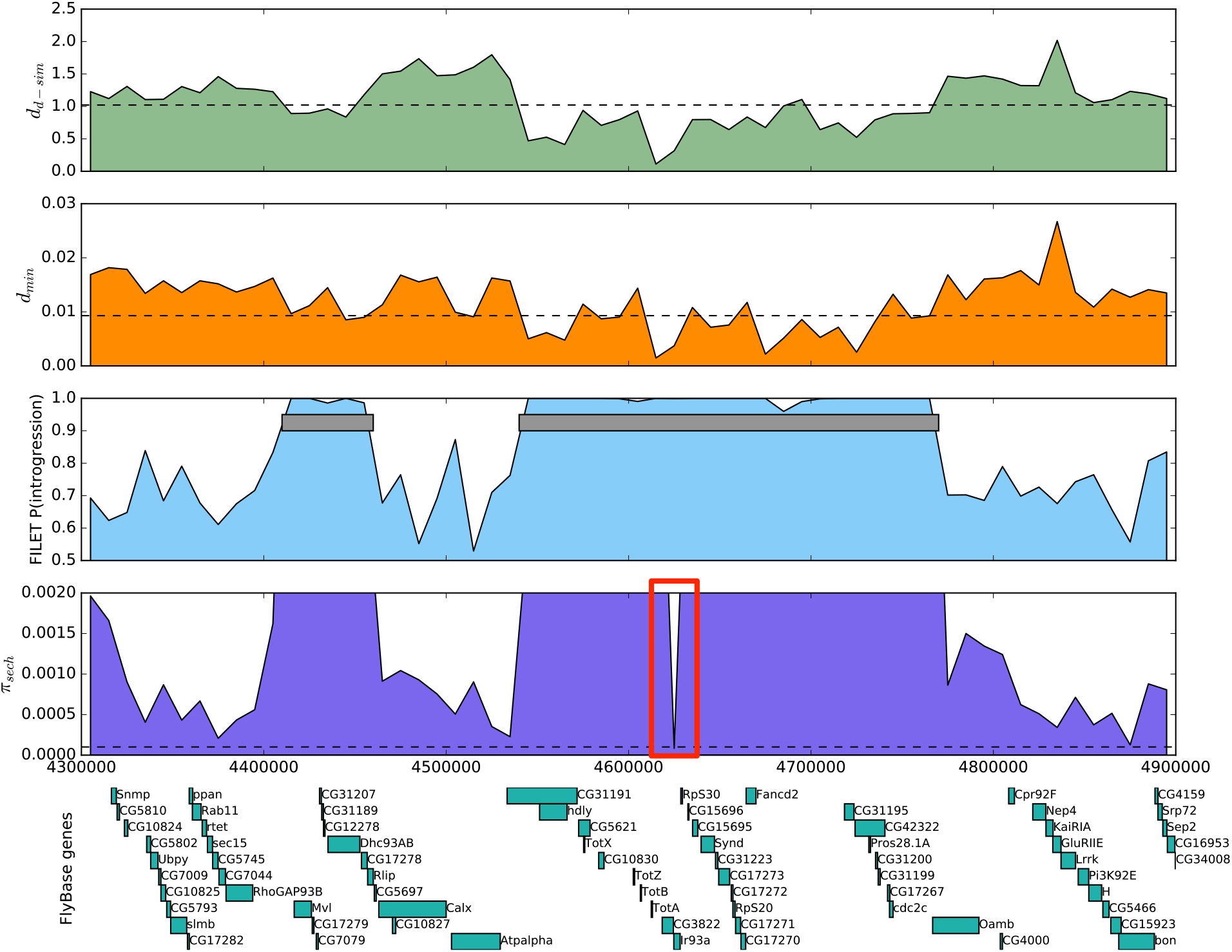
A large genomic region on 3R classified by FILET as introgressed from *D. simulans* to *D. sechellia*. Values of the *d_d_*_-*sim*_ and *d_min_* (upper two panels) within each 10 kb window in the region are shown, along with the posterior probability of introgression from FILET (i.e. 1 – *P*(no introgression)). Clustered regions classified as introgressed are shown as gray rectangles superimposed over these probabilities. Also shown are windowed values of *π* in *D. sechellia*, with the sweep region highlighted in red, and the locations of annotated genes with associated FlyBase identifiers [90].

Interestingly, this particular window is the only one in this region that is classified by FILET as having recent gene flow from *D. sechellia* to *D. simulans*. However this classification may be erroneous as one might expect FILET, which was not trained on any examples of adaptive introgression, to make an error in such a scenario because rather than gene flow increasing polymorphism in the recipient population, diversity is greatly diminished if the introgressed alleles rapidly sweep toward fixation. We note that this window is immediately flanked by a large number of windows classified as introgressed from *D. simulans* to *D. sechellia* and which show a large increase in diversity in the recipient population as expected. Moreover, Brand et al.’s phylogenetic analysis of introgression in this region also supported gene flow in this direction. Brand et al. also found evidence suggesting that the introgressed haplotype began sweeping to higher frequency in *D. simulans* (though it has not reached fixation in this species) prior to the timing of the introgression and subsequent sweep in *D. sechellia*. Thus we conclude that the adaptive allele probably did indeed originate in *D. simulans* before migrating to *D. sechellia*, and FILET’s apparent error in this case underscores the genealogical differences between adaptive gene flow and introgression events involving only neutral alleles.

### Concluding remarks

Here we present a novel machine learning approach, FILET, that leverages population genomic data from two related populations in order to determine whether a given genomic window has experienced gene flow between these populations, and if so in which direction. We applied FILET to a set of *D. simulans* genomes as well as a new set of whole genome sequences from the closely related island endemic *D. sechellia*, confirming widespread introgression and also inferring that this introgression was largely in the direction of *D. simulans* to *D. sechellia*. Future work leveraging *D. simulans* data sampled from the Seychelles will be required to determine whether this asymmetry is a consequence of low rate of migration of *D. simulans* back to mainland Africa (where our *D. simulans* data were obtained), or whether the directionality of gene flow is biased on the islands themselves. In addition to creating FILET, we devised several new statistics, including the *d_d_* statistics and *Z_X_* which our feature rankings show to be quite useful for uncovering gene flow.

Despite the success of FILET on both simulated data sets and real data from *Drosophila*, there are several improvements that could be made. First, by framing the problem as one of parameter estimation (i.e. regression) rather than classification, we may be able to precisely infer the values of relevant parameters of introgression events (i.e. the time of the event and the number of migrant lineages). Deep learning methods, which naturally allow for both classification and regression, may prove particularly useful for this task [103]. Indeed, Sheehan and Song [109] used deep learning to infer demographic parameters (regression) while simultaneously identifying selective sweeps (classification). Another step we have not taken is to explicitly handle adaptive introgression, which could potentially greatly improve our approach’s power to detect such events.

While population genetic inference has traditionally relied on the design of a summary statistic sensitive to the evolutionary force of interest, a number of highly successful supervised machine learning methods have been put forth within the last few years [54, 104–109]. These methods are often thought of as black boxes, a characterization that may not always be fair [125]. Indeed in the context of evolutionary genetics such machine learning approaches are easily interpreted as we have strong generative models that guide our intuition. Nonetheless, classical statistical estimation from parametric models may often be more interpretable. Hybrid approaches combining machine learning techniques with Bayesian approaches to estimate posterior distributions of evolutionary parameters (e.g. [127]) thus represent an attractive alternative to either approach in their “pure” form. As genomic data sets continue to grow, we argue that machine learning approaches—in whatever shape they eventually take—leveraging high dimensional feature spaces have the potential to revolutionize evolutionary genomic inference.

## ACKNOWLEDGMENTS

We thank Michael Lan for his work on an early iteration of this project. We also thank Joshua Schraiber and two anonymous reviewers for comments on the manuscript.

## SUPPLEMENTAL FIGURE AND TABLE LEGENDS

**S1 Fig.** Illustration of the difference in values of the *d_min_* statistic calculated from joint population samples with and without introgression.

**S2 Fig.** Violin plots showing the values of *d_min_*, all four *d_d_* statistics, and *Z_X_* under simulated scenarios including introgression or lacking it for each values of *T_D_*. The values of these statistics were obtained from the training data sets described in the Materials and Methods.

**S3 Fig.** Heatmaps showing several methods’ sensitivity to detect introgression. Same as Figure 1, but for other values of *T_D_*. (*A*) Accuracy for *d_min_* and *G_min_* when *T_D_*= 1×4*N* generations. (*B*) Accuracy of FILET when *T_D_* = 1×4*N*. (*C*) and (*D*) show the same when *T_D_* = 4×4*N*. (E) and (F) show the same when 16×4*N*.

**S4 Fig.** Heatmaps showing FILET’s sensitivity to introgression from an unsampled ghost population. (*A*) Sensitivity when *T_D_*= 0.25×4*N* generations. (*B*) Sensitivity when *T_D_* = 1×4*N* generations. (*C*) *T_D_* = 4×4*N* generations. (*D*) *T_D_* = 16×4*N*.

**S5 Fig.** ROC curves showing power of FILET, *d_min_* and *G_min_* under each value of *T_D_*. In order to generate these curves we transformed the classification task into a binary one: discriminating between isolation and introgression in either direction. (*A*) *T_D_* = 0.25×4*N* generations. (*B*) *T_D_* = 1×4*N* generations. (*C*) *T_D_* = 4×4*N*. (*D*) *T_D_* = 16×4*N*. Training and test sets for these problems contained equal numbers of examples of introgression from population 1 into 2 and introgression from population 2 into 1.

**S6 Fig.** ROC curves showing power of versions of FILET trained with decreasing numbers of training instances (ranging from 100 to 10000 for each class). (*A*) *T_D_* = 0.25×4*N* generations. (*B*) *T_D_* = 1×4*N* generations. (*C*) *T_D_*= 4×4*N*. (*D*) *T_D_* = 16×4*N*.

**S7 Fig.** ROC curve showing FILET’s power when trained and tested on simulated examples with different window sizes with *T_D_* = 0.25×4*N* generations.

**S8 Fig.** A comparison of the positive predictive value of FILET and ChromoPainter on the same simulated data used for Fig 2. The “coarse” and “fine” versions of FILET, and lenient and stringent versions of ChromoPainter’s predictions, are as defined for Fig 2. In cases where the positive predictive value is undefined (i.e. no base pairs were predicted to be introgressed), it is displayed as zero (i.e. a white cell in the heatmap).

**S9 Fig.** Population structure within *D. sechellia.* (*A*) The top three principal components of all *D. sechellia* diploid genomes. The cluster on the left shows the individuals from Praslin, while the cluster on the right shows all other individuals. Note that the cluster on the right is far less dispersed due to the very small amount of polymorphism among these individuals. The numbers in parentheses on each axis show the fraction of the variance explained by each principal component. (*B*) Results of running fastStructure on our *D. sechellia* samples with the number of subpopulations (*K*) ranging from 2 to 8.

**S10 Fig.** Site frequency spectra of *D. sechellia* samples from Praslin, *D. sechellia* samples from all other locations, and *D. simulans* samples. The *D. sechellia* samples were both downsampled to *n*=12 as described in the text, while *D. simulans* was downsampled to *n*=18 (i.e. the same sample sizes used for our demographic inference). These SFS show the fraction of all polymorphisms found in each bin rather than the raw number of polymorphisms, and thus do not contain information about the total number of SNPs. As described in the text, there is >12-fold more polymorphism in the Praslin samples than in the non-Praslin samples.

**S11 Fig.** Confusion matrix showing FILET’s classification accuracy when trained under out inferred model of the *simulans*-*sechellia* joint demographic history, but applied to test data generated under a different model (described in Materials and Methods and shown in S4 Table). under this model as assessed on an independent simulated test set. Perfect accuracy would be 100% along the entire diagonal from top-left to bottom-right, and the false positive rate is the sum of top-middle and top-right cells.

**S1 Table.** Results when applying FILET to simulations with constant bidirectional migration. 10000 simulated replicates were tested for each parameter combination.

**S2 Table.** Feature importance and rankings for each classifier used in this study.

**S3 Table.** Sampling location, sequencing/mapping statistics, and SRA identifiers for each genome included in this study.

**S4 Table.** Demographic parameter estimates inferred by *∂*a*∂*i, along with a simple naïve model.

## REFERENCES

1. Mallet J. Hybridization as an invasion of the genome. Trends in ecology & evolution. 2005;20(5):229–37.

2. Whitney KD, Ahern JR, Campbell LG, Albert LP, King MS. Patterns of hybridization in plants. Perspectives in Plant Ecology, Evolution and Systematics. 2010;12(3):175–82.

3. Barton NH. The role of hybridization in evolution. Mol Ecol. 2001;10(3):551–68.

4. Tung J, Barreiro LB. The contribution of admixture to primate evolution. Current opinion in genetics & development. 2017;47:61–8.

5. Baack EJ, Rieseberg LH. A genomic view of introgression and hybrid speciation. Current opinion in genetics & development. 2007;17(6):513–8.

6. Goulet BE, Roda F, Hopkins R. Hybridization in plants: old ideas, new techniques. Plant Physiol. 2017;173(1):65–78.

7. Gladieux P, Ropars J, Badouin H, Branca A, Aguileta G, Vienne DM, et al. Fungal evolutionary genomics provides insight into the mechanisms of adaptive divergence in eukaryotes. Mol Ecol. 2014;23(4):753–73.

8. Schardl C, Craven K. Interspecific hybridization in plant-associated fungi and oomycetes: a review. Mol Ecol. 2003;12(11):2861–73.

9. Brandvain Y, Kenney AM, Flagel L, Coop G, Sweigart AL. Speciation and introgression between Mimulus nasutus and Mimulus guttatus. PLoS Genet. 2014;10(6):e1004410.

10. Begun DJ, Holloway AK, Stevens K, Hillier LW, Poh Y-P, Hahn MW, et al. Population genomics: whole-genome analysis of polymorphism and divergence in Drosophila simulans. PLoS Biol. 2007;5(11):e310.

11. Kulathinal RJ, Stevison LS, Noor MA. The genomics of speciation in Drosophila: diversity, divergence, and introgression estimated using low-coverage genome sequencing. PLoS Genet. 2009;5(7):e1000550.

12. Martin SH, Dasmahapatra KK, Nadeau NJ, Salazar C, Walters JR, Simpson F, et al. Genome-wide evidence for speciation with gene flow in Heliconius butterflies. Genome Res. 2013;23(11):1817–28.

13. Fontaine MC, Pease JB, Steele A, Waterhouse RM, Neafsey DE, Sharakhov IV, et al. Extensive introgression in a malaria vector species complex revealed by phylogenomics. Science. 2015;347(6217):1258524.

14. Nürnberger B, Lohse K, Fijarczyk A, Szymura JM, Blaxter ML. Para-allopatry in hybridizing fire-bellied toads (Bombina bombina and B. variegata): Inference from transcriptome-wide coalescence analyses. Evolution. 2016;70(8):1803–18.

15. Rothfels CJ, Johnson AK, Hovenkamp PH, Swofford DL, Roskam HC, Fraser-Jenkins CR, et al. Natural hybridization between genera that diverged from each other approximately 60 million years ago. The American Naturalist. 2015;185(3):433–42.

16. Nadeau NJ, Ruiz M, Salazar P, Counterman B, Medina JA, Ortiz-Zuazaga H, et al. Population genomics of parallel hybrid zones in the mimetic butterflies, H. melpomene and H. erato. Genome Res. 2014;24(8):1316–33.

17. Turissini DA, Matute DR. Fine scale mapping of genomic introgressions within the Drosophila yakuba clade. bioRxiv. 2017:152421.

18. Bachtrog D, Thornton K, Clark A, Andolfatto P. Extensive introgression of mitochondrial DNA relative to nuclear genes in the Drosophila yakuba species group. Evolution. 2006;60(2):292–302.

19. Leavitt DH, Marion AB, Hollingsworth BD, Reeder TW. Multilocus phylogeny of alligator lizards (Elgaria, Anguidae): Testing mtDNA introgression as the source of discordant molecular phylogenetic hypotheses. Mol Phylogenet Evol. 2017;110:104–21.

20. Sarver BA, Demboski JR, Good JM, Forshee N, Hunter SS, Sullivan J. Comparative phylogenomic assessment of mitochondrial introgression among several species of chipmunks (Tamias). Genome Biol Evol. 2016;9(1):7–19.

21. Carneiro M, Albert FW, Afonso S, Pereira RJ, Burbano H, Campos R, et al. The genomic architecture of population divergence between subspecies of the European rabbit. PLoS Genet. 2014;10(8):e1003519.

22. Maroja LS, Larson EL, Bogdanowicz SM, Harrison RG. Genes with restricted introgression in a field cricket (Gryllus firmus/Gryllus pennsylvanicus) hybrid zone are concentrated on the X chromosome and a single autosome. G3: Genes, Genomes, Genetics. 2015;5(11):2219-27.

23. Muirhead CA, Presgraves DC. Hybrid incompatibilities, local adaptation, and the genomic distribution of natural introgression between species. The American Naturalist. 2016;187(2):249–61.

24. Phifer-Rixey M, Bomhoff M, Nachman MW. Genome-wide patterns of differentiation among house mouse subspecies. Genetics. 2014;198(1):283–97.

25. Green RE, Krause J, Briggs AW, Maricic T, Stenzel U, Kircher M, et al. A draft sequence of the Neandertal genome. Science. 2010;328(5979):710-22.

26. Sankararaman S, Mallick S, Dannemann M, Prüfer K, Kelso J, Pääbo S, et al. The genomic landscape of Neanderthal ancestry in present-day humans. Nature. 2014;507(7492):354-7.

27. Turner TL, Hahn MW, Nuzhdin SV. Genomic islands of speciation in Anopheles gambiae. PLoS Biol. 2005;3(9):e285.

28. Harris K, Nielsen R. The genetic cost of Neanderthal introgression. Genetics. 2016;203(2):881–91.

29. Juric I, Aeschbacher S, Coop G. The strength of selection against Neanderthal introgression. PLoS Genet. 2016;12(11):e1006340.

30. Hedrick PW. Adaptive introgression in animals: examples and comparison to new mutation and standing variation as sources of adaptive variation. Mol Ecol. 2013;22(18):4606–18.

31. Norris LC, Main BJ, Lee Y, Collier TC, Fofana A, Cornel AJ, et al. Adaptive introgression in an African malaria mosquito coincident with the increased usage of insecticide-treated bed nets. Proceedings of the National Academy of Sciences. 2015;112(3):815–20.

32. Pardo-Diaz C, Salazar C, Baxter SW, Merot C, Figueiredo-Ready W, Joron M, et al. Adaptive introgression across species boundaries in Heliconius butterflies. PLoS Genet. 2012;8(6):e1002752.

33. Song Y, Endepols S, Klemann N, Richter D, Matuschka F-R, Shih C-H, et al. Adaptive introgression of anticoagulant rodent poison resistance by hybridization between old world mice. Curr Biol. 2011;21(15):1296–301.

34. Bechsgaard J, Jorgensen TH, Schierup MH. Evidence for Adaptive Introgression of Disease Resistance Genes Among Closely Related Arabidopsis Species. G3: Genes, Genomes, Genetics. 2017;7(8):2677-83.

35. Cheeseman K, Ropars J, Renault P, Dupont J, Gouzy J, Branca A, et al. Multiple recent horizontal transfers of a large genomic region in cheese making fungi. Nature Communications. 2014;5:2876.

36. Huerta-Sánchez E, Jin X, Bianba Z, Peter BM, Vinckenbosch N, Liang Y, et al. Altitude adaptation in Tibetans caused by introgression of Denisovan-like DNA. Nature. 2014;512(7513):194-7.

37. Melo MC, Salazar C, Jiggins CD, Linares M. Assortative mating preferences among hybrids offers a route to hybrid speciation. Evolution. 2009;63(6):1660–5.

38. Salazar C, Baxter SW, Pardo-Diaz C, Wu G, Surridge A, Linares M, et al. Genetic evidence for hybrid trait speciation in Heliconius butterflies. PLoS Genet. 2010;6(4):e1000930.

39. Alexander DH, Novembre J, Lange K. Fast model-based estimation of ancestry in unrelated individuals. Genome Res. 2009;19(9):1655–64.

40. Anderson E, Thompson E. A model-based method for identifying species hybrids using multilocus genetic data. Genetics. 2002;160(3):1217–29.

41. Pritchard JK, Stephens M, Donnelly P. Inference of population structure using multilocus genotype data. Genetics. 2000;155(2):945–59.

42. Pickrell JK, Pritchard JK. Inference of population splits and mixtures from genome-wide allele frequency data. PLoS Genet. 2012;8(11):e1002967.

43. Guan Y. Detecting structure of haplotypes and local ancestry. Genetics. 2014;196(3):625–42.

44. Tang H, Coram M, Wang P, Zhu X, Risch N. Reconstructing genetic ancestry blocks in admixed individuals. The American Journal of Human Genetics. 2006;79(1):1–12.

45. Sohn K-A, Ghahramani Z, Xing EP. Robust estimation of local genetic ancestry in admixed populations using a nonparametric Bayesian approach. Genetics. 2012;191(4):1295–308.

46. Lawson DJ, Hellenthal G, Myers S, Falush D. Inference of population structure using dense haplotype data. PLoS Genet. 2012;8(1):e1002453.

47. Price AL, Tandon A, Patterson N, Barnes KC, Rafaels N, Ruczinski I, et al. Sensitive detection of chromosomal segments of distinct ancestry in admixed populations. PLoS Genet. 2009;5(6):e1000519.

48. Wright S. The genetical structure of populations. Ann Hum Genet. 1949;15(1):323–54.

49. Nei M, Li W-H. Mathematical model for studying genetic variation in terms of restriction endonucleases. Proceedings of the National Academy of Sciences. 1979;76(10):5269–73.

50. Joly S, McLenachan PA, Lockhart PJ. A statistical approach for distinguishing hybridization and incomplete lineage sorting. The American Naturalist. 2009;174(2):E54–E70.

51. Geneva AJ, Muirhead CA, Kingan SB, Garrigan D. A new method to scan genomes for introgression in a secondary contact model. PLoS ONE. 2015;10(4):e0118621.

52. Rosenzweig BK, Pease JB, Besansky NJ, Hahn MW. Powerful methods for detecting introgressed regions from population genomic data. Mol Ecol. 2016;25(11):2387–97. doi: 10.1111/mec.13610.

53. Geurts P, Ernst D, Wehenkel L. Extremely randomized trees. Machine Learning. 2006;63(1):3–42.

54. Schrider DR, Kern AD. S/HIC: Robust Identification of Soft and Hard Sweeps Using Machine Learning. PLoS Genet. 2016;12(3): e1005928.

55. Schrider DR, Kern AD. Soft sweeps are the dominant mode of adaptation in the human genome. Mol Biol Evol. 2017;34(8):1863–77. Epub 2017/05/10. doi: 10.1093/molbev/msx154. PubMed PMID: 28482049.

56. Jones CD. The genetic basis of Drosophila sechellia’s resistance to a host plant toxin. Genetics. 1998;149(4):1899–908.

57. Jones CD. The genetics of adaptation in Drosophila sechellia. Genetica. 2005;123(1-2):137.

58. Louis J, David J. Ecological specialization in the Drosophila melanogaster species subgroup: a case study of D. sechellia. Acta oecologica Oecologia generalis. 1986;7(3):215–29.

59. Farine J-P, Legal L, Moreteau B, Le Quere J-L. Volatile components of ripe fruits of Morinda citrifolia and their effects on Drosophila. Phytochemistry. 1996;41(2):433–8.

60. Legal L, Chappe B, Jallon JM. Molecular basis ofMorinda citrifolia (L.): Toxicity on drosophila. J Chem Ecol. 1994;20(8):1931–43.

61. Legal L, Moulin B, Jallon JM. The relation between structures and toxicity of oxygenated aliphatic compounds homologous to the insecticide octanoic acid and the chemotaxis of two species of Drosophila. Pestic Biochem Physiol. 1999;65(2):90–101.

62. Andrade López J, Lanno S, Auerbach J, Moskowitz E, Sligar L, Wittkopp P, et al. Genetic basis of octanoic acid resistance in Drosophila sechellia: functional analysis of a fine-mapped region. Mol Ecol. 2017;26(4):1148–60.

63. Dekker T, Ibba I, Siju K, Stensmyr MC, Hansson BS. Olfactory shifts parallel superspecialism for toxic fruit in Drosophila melanogaster sibling, D. sechellia. Curr Biol. 2006;16(1):101–9.

64. Huang Y, Erezyilmaz D. The genetics of resistance to Morinda fruit toxin during the postembryonic stages in Drosophila sechellia. G3: Genes, Genomes, Genetics. 2015;5(10):1973-81.

65. Hungate EA, Earley EJ, Boussy IA, Turissini DA, Ting C-T, Moran JR, et al. A locus in Drosophila sechellia affecting tolerance of a host plant toxin. Genetics. 2013;195(3):1063–75.

66. Matsuo T, Sugaya S, Yasukawa J, Aigaki T, Fuyama Y. Odorant-binding proteins OBP57d and OBP57e affect taste perception and host-plant preference in Drosophila sechellia. PLoS Biol. 2007;5(5):e118.

67. Shiao M-S, Chang J-M, Fan W-L, Lu M-YJ, Notredame C, Fang S, et al. Expression divergence of chemosensory genes between Drosophila sechellia and its sibling species and its implications for host shift. Genome Biol Evol. 2015;7(10):2843–58.

68. Hey J, Kliman RM. Population genetics and phylogenetics of DNA sequence variation at multiple loci within the Drosophila melanogaster species complex. Mol Biol Evol. 1993;10(4):804–22.

69. Kern AD, Jones CD, Begun DJ. Molecular population genetics of male accessory gland proteins in the Drosophila simulans complex. Genetics. 2004;167(2):725–35.

70. Kliman RM, Andolfatto P, Coyne JA, Depaulis F, Kreitman M, Berry AJ, et al. The population genetics of the origin and divergence of the Drosophila simulans complex species. Genetics. 2000;156(4):1913–31.

71. Legrand D, Tenaillon MI, Matyot P, Gerlach J, Lachaise D, Cariou M-L. Species-wide genetic variation and demographic history of Drosophila sechellia, a species lacking population structure. Genetics. 2009;182(4):1197–206.

72. Garrigan D, Kingan SB, Geneva AJ, Andolfatto P, Clark AG, Thornton KR, et al. Genome sequencing reveals complex speciation in the Drosophila simulans clade. Genome Res. 2012;22(8):1499–511.

73. Matute D, Ayroles J. Hybridization occurs between Drosophila simulans and D. sechellia in the Seychelles archipelago. J Evol Biol. 2014;27(6):1057–68.

74. Rogers RL, Cridland JM, Shao L, Hu TT, Andolfatto P, Thornton KR. Landscape of standing variation for tandem duplications in Drosophila yakuba and Drosophila simulans. Mol Biol Evol. 2014;31(7):1750–66.

75. Feder JL, Xie X, Rull J, Velez S, Forbes A, Leung B, et al. Mayr, Dobzhansky, and Bush and the complexities of sympatric speciation in Rhagoletis. Proceedings of the National Academy of Sciences. 2005;102(suppl 1):6573–80.

76. Kelly JK. A test of neutrality based on interlocus associations. Genetics. 1997;146(3):1197–206.

77. Breiman L. Random forests. Machine Learning. 2001;45(1):5–32.

78. Quinlan JR. Induction of decision trees. Machine Learning. 1986;1(1):81–106.

79. Fay JC, Wu C-I. Hitchhiking under positive Darwinian selection. Genetics. 2000;155(3):1405–13.

80. Tajima F. Statistical method for testing the neutral mutation hypothesis by DNA polymorphism. Genetics. 1989;123(3):585–95.

81. Hudson RR, Slatkin M, Maddison W. Estimation of levels of gene flow from DNA sequence data. Genetics. 1992;132(2):583–9.

82. Hudson RR. A new statistic for detecting genetic differentiation. Genetics. 2000;155(4):2011–4.

83. Patterson N, Moorjani P, Luo Y, Mallick S, Rohland N, Zhan Y, et al. Ancient admixture in human history. Genetics. 2012;192(3):1065–93.

84. Pedregosa F, Varoquaux G, Gramfort A, Michel V, Thirion B, Grisel O, et al. Scikit-learn: Machine learning in Python. Journal of Machine Learning Research. 2011;12(Oct):2825–30.

85. Breiman L, Friedman J, Stone CJ, Olshen RA. Classification and regression trees: CRC press; 1984.

86. Chan AH, Jenkins PA, Song YS. Genome-wide fine-scale recombination rate variation in Drosophila melanogaster. PLoS Genet. 2012;8(12):e1003090.

87. Li N, Stephens M. Modeling linkage disequilibrium and identifying recombination hotspots using single-nucleotide polymorphism data. Genetics. 2003;165(4):2213–33.

88. Li H. Aligning sequence reads, clone sequences and assembly contigs with BWA-MEM. arXiv. 2013. doi: 1303.3997.

89. Hu TT, Eisen MB, Thornton KR, Andolfatto P. A second-generation assembly of the *Drosophila simulans* genome provides new insights into patterns of lineage-specific divergence. Genome Res. 2013;23(1):89–98.

90. Gramates LS, Marygold SJ, Santos Gd, Urbano J-M, Antonazzo G, Matthews BB, et al. FlyBase at 25: looking to the future. Nucleic Acids Res. 2017;45(D1):D663–D71.

91. McKenna A, Hanna M, Banks E, Sivachenko A, Cibulskis K, Kernytsky A, et al. The Genome Analysis Toolkit: a MapReduce framework for analyzing next-generation DNA sequencing data. Genome Res. 2010;20(9):1297–303.

92. DePristo MA, Banks E, Poplin R, Garimella KV, Maguire JR, Hartl C, et al. A framework for variation discovery and genotyping using next-generation DNA sequencing data. Nat Genet. 2011;43(5):491–8.

93. Auwera GA, Carneiro MO, Hartl C, Poplin R, del Angel G, Levy-Moonshine A, et al. From FastQ data to high-confidence variant calls: the genome analysis toolkit best practices pipeline. Current protocols in bioinformatics. 2013;43:11.0. 1-.0. 33.

94. Delaneau O, Zagury J-F, Marchini J. Improved whole-chromosome phasing for disease and population genetic studies. Nat Methods. 2013;10(1):5–6.

95. Gutenkunst RN, Hernandez RD, Williamson SH, Bustamante CD. Inferring the joint demographic history of multiple populations from multidimensional SNP frequency data. PLoS Genet. 2009;5(10):e1000695.

96. Jansen PW, Perez RE. Constrained structural design optimization via a parallel augmented Lagrangian particle swarm optimization approach. Computers & Structures. 2011;89(13):1352–66.

97. Kraft D. A software package for sequential quadratic programming: DFVLR Obersfaffeuhofen, Germany; 1988.

98. Perez RE, Jansen PW, Martins JR. pyOpt: a Python-based object-oriented framework for nonlinear constrained optimization. Structural and Multidisciplinary Optimization. 2012;45(1):101–18.

99. Schrider DR, Shanku AG, Kern AD. Effects of Linked Selective Sweeps on Demographic Inference and Model Selection. Genetics. 2016;204(3):1207–23. doi: 10.1534/genetics.116.190223.

100. Pool JE. The mosaic ancestry of the Drosophila genetic reference panel and the D. melanogaster reference genome reveals a network of epistatic fitness interactions. Mol Biol Evol. 2015;32(12):3236–51.

101. Schrider DR, Kern AD. Supervised Machine Learning for Population Genetics: A New Paradigm. Trends Genet. 2018:10.1016/j.tig.2017.12.005.

102. Cortes C, Vapnik V. Support-vector networks. Machine Learning. 1995;20(3):273–97.

103. LeCun Y, Bengio Y, Hinton G. Deep learning. Nature. 2015;521(7553):436-44.

104. Lin K, Li H, Schlötterer C, Futschik A. Distinguishing positive selection from neutral evolution: boosting the performance of summary statistics. Genetics. 2011;187(1):229–44.

105. Pavlidis P, Jensen JD, Stephan W. Searching for footprints of positive selection in whole-genome SNP data from nonequilibrium populations. Genetics. 2010;185(3):907–22.

106. Pybus M, Luisi P, Dall’Olio GM, Uzkudun M, Laayouni H, Bertranpetit J, et al. Hierarchical boosting: a machine-learning framework to detect and classify hard selective sweeps in human populations. Bioinformatics. 2015;31(24):3946–52.

107. Ronen R, Udpa N, Halperin E, Bafna V. Learning natural selection from the site frequency spectrum. Genetics. 2013;195(1):181–93.

108. Pudlo P, Marin J-M, Estoup A, Cornuet J-M, Gautier M, Robert CP. Reliable ABC model choice via random forests. Bioinformatics. 2016;32(6):859–66.

109. Sheehan S, Song YS. Deep learning for population genetic inference. PLoS Comput Biol. 2016;12(5):e1004845.

110. Gazave E, Ma L, Chang D, Coventry A, Gao F, Muzny D, et al. Neutral genomic regions refine models of recent rapid human population growth. Proceedings of the National Academy of Sciences. 2014;111(2):757–62.

111. Ewing GB, Jensen JD. The consequences of not accounting for background selection in demographic inference. Mol Ecol. 2016;25(1):135–41.

112. Schrider DR, Houle D, Lynch M, Hahn MW. Rates and genomic consequences of spontaneous mutational events in Drosophila melanogaster. Genetics. 2013;194(4):937–54.

113. Langley CH, Stevens K, Cardeno C, Lee YCG, Schrider DR, Pool JE, et al. Genomic variation in natural populations of *Drosophila melanogaster*. Genetics. 2012;192(2):533–98.

114. Raj A, Stephens M, Pritchard JK. fastSTRUCTURE: variational inference of population structure in large SNP data sets. Genetics. 2014;197(2):573–89.

115. Legrand D, Vautrin D, Lachaise D, Cariou M-L. Microsatellite variation suggests a recent fine-scale population structure of Drosophila sechellia, a species endemic of the Seychelles archipelago. Genetica. 2011;139(7):909.

116. Navascués M, Legrand D, Campagne C, Cariou M-L, Depaulis F. Distinguishing migration from isolation using genes with intragenic recombination: detecting introgression in the Drosophila simulans species complex. BMC Evol Biol. 2014;14(1):89.

117. Obbard DJ, Maclennan J, Kim K-W, Rambaut A, O’Grady PM, Jiggins FM. Estimating divergence dates and substitution rates in the Drosophila phylogeny. Mol Biol Evol. 2012;29(11):3459–73.

118. Keightley PD, Trivedi U, Thomson M, Oliver F, Kumar S, Blaxter M. Analysis of the genome sequences of three Drosophila melanogaster spontaneous mutation accumulation lines. Genome Res. 2009;19:1195–201.

119. Brand CL, Kingan SB, Wu L, Garrigan D. A selective sweep across species boundaries in Drosophila. Mol Biol Evol. 2013;30(9):2177–86.

120. Benton R, Vannice KS, Gomez-Diaz C, Vosshall LB. Variant ionotropic glutamate receptors as chemosensory receptors in Drosophila. Cell. 2009;136(1):149–62.

121. Lu H-L, Wang JB, Brown MA, Euerle C, Leger RJS. Identification of Drosophila mutants affecting defense to an entomopathogenic fungus. Scientific reports. 2015;5.

122. Ekengren S, Hultmark D. A family of Turandot-related genes in the humoral stress response of Drosophila. Biochem Biophys Res Commun. 2001;284(4):998–1003.

123. Ekengren S, Tryselius Y, Dushay MS, Liu G, Steiner H, Hultmark D. A humoral stress response in Drosophila. Curr Biol. 2001;11(9):714–8.

124. Salazar-Jaramillo L, Jalvingh KM, de Haan A, Kraaijeveld K, Buermans H, Wertheim B. Inter-and intra-species variation in genome-wide gene expression of Drosophila in response to parasitoid wasp attack. BMC Genomics. 2017;18(1):331.

125. Breiman L. Statistical modeling: The two cultures (with comments and a rejoinder by the author). Statistical science. 2001;16(3):199–231.

126. Baehrens D, Schroeter T, Harmeling S, Kawanabe M, Hansen K, MÃžller K-R. How to explain individual classification decisions. Journal of Machine Learning Research. 2010;11(Jun):1803–31.

127. Blum MG, François O. Non-linear regression models for Approximate Bayesian Computation. Statistics and Computing. 2010;20(1):63–73.

