## Supplementary figures and images for "Supervised machine learning reveals introgressed loci in the genomes of *Drosophila simulans* and *D. sechellia*"

### S1_Fig.pdf

Figure S1

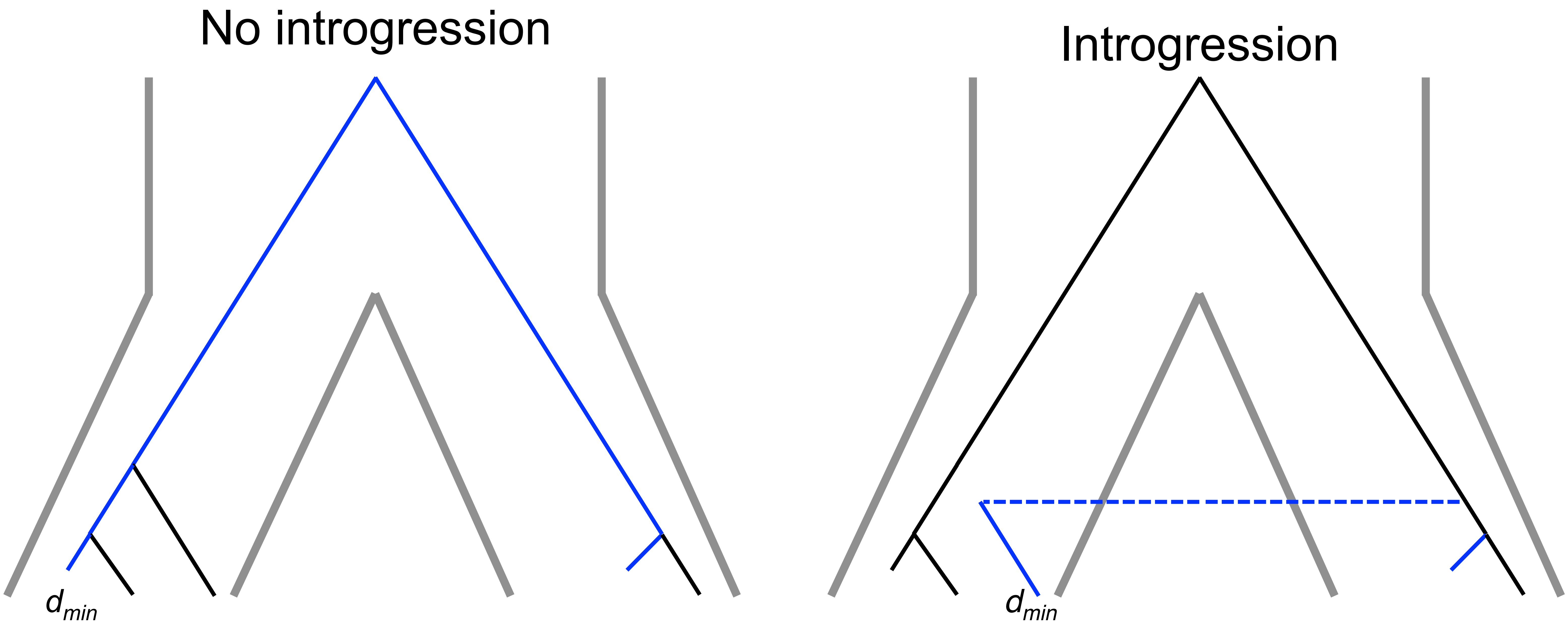

### S2_Fig.pdf

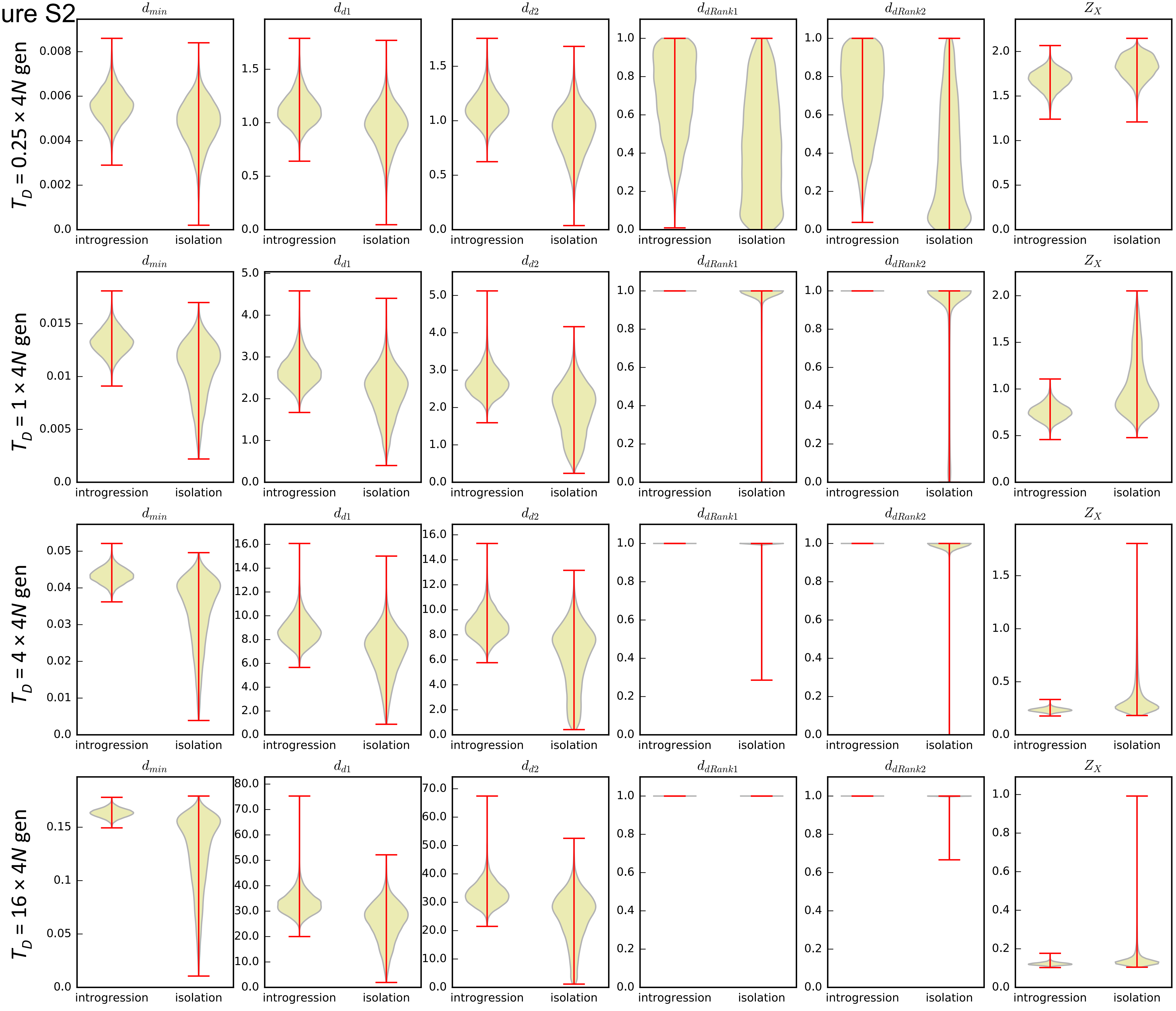

### S3_Fig.pdf

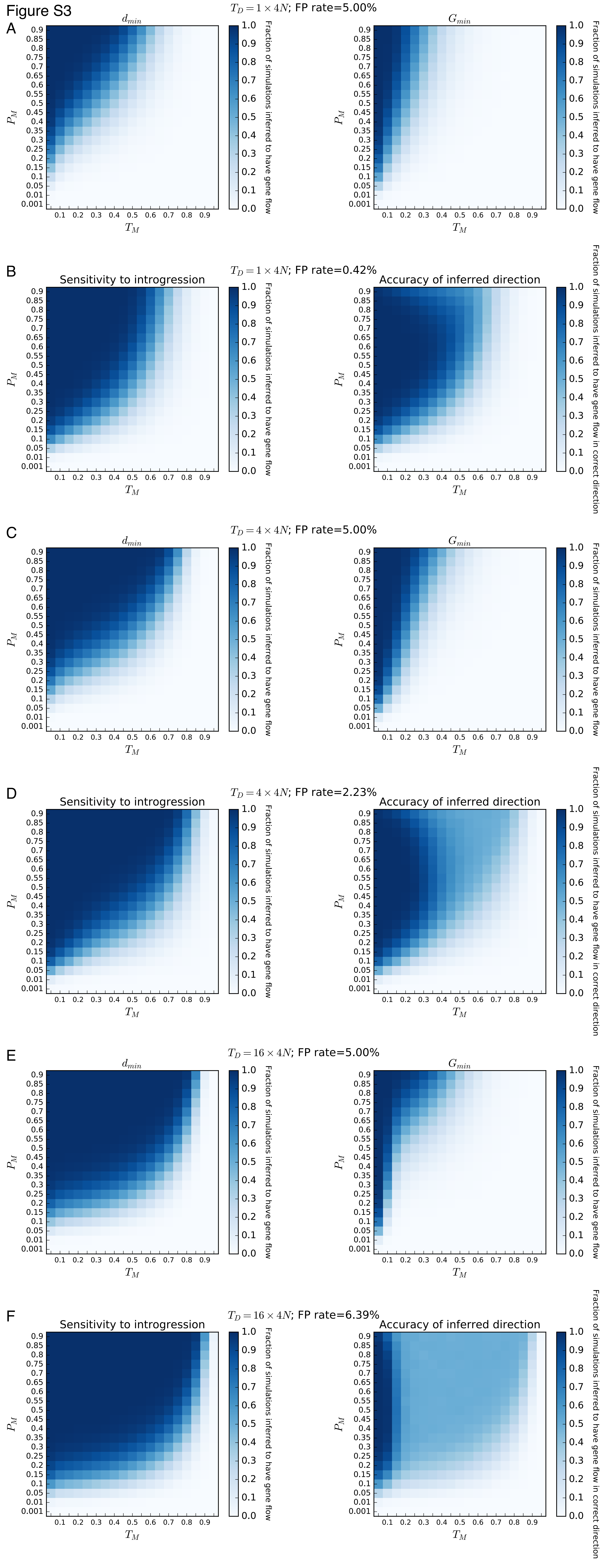

### S4_Fig.pdf

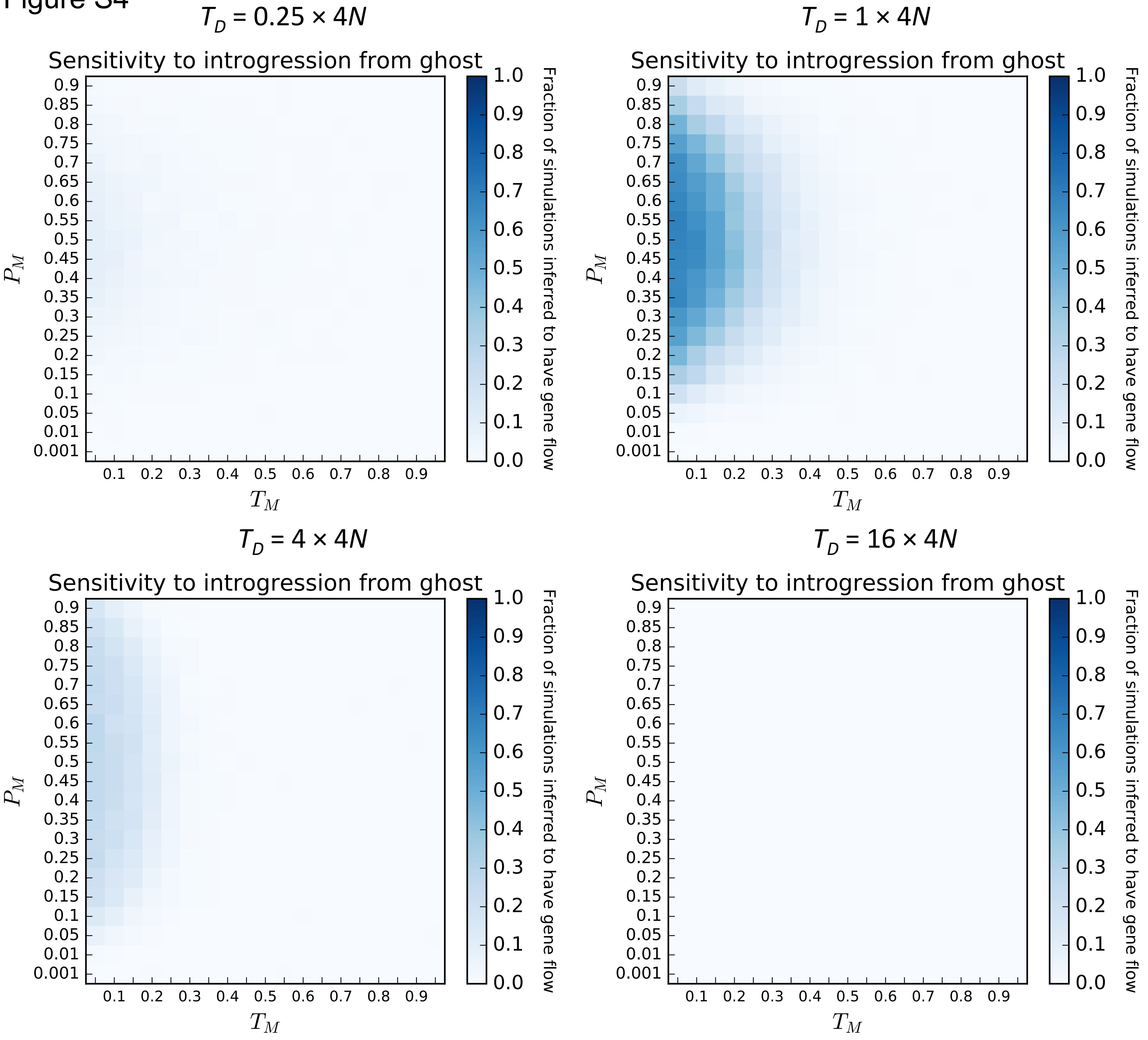

### S5_Fig.pdf

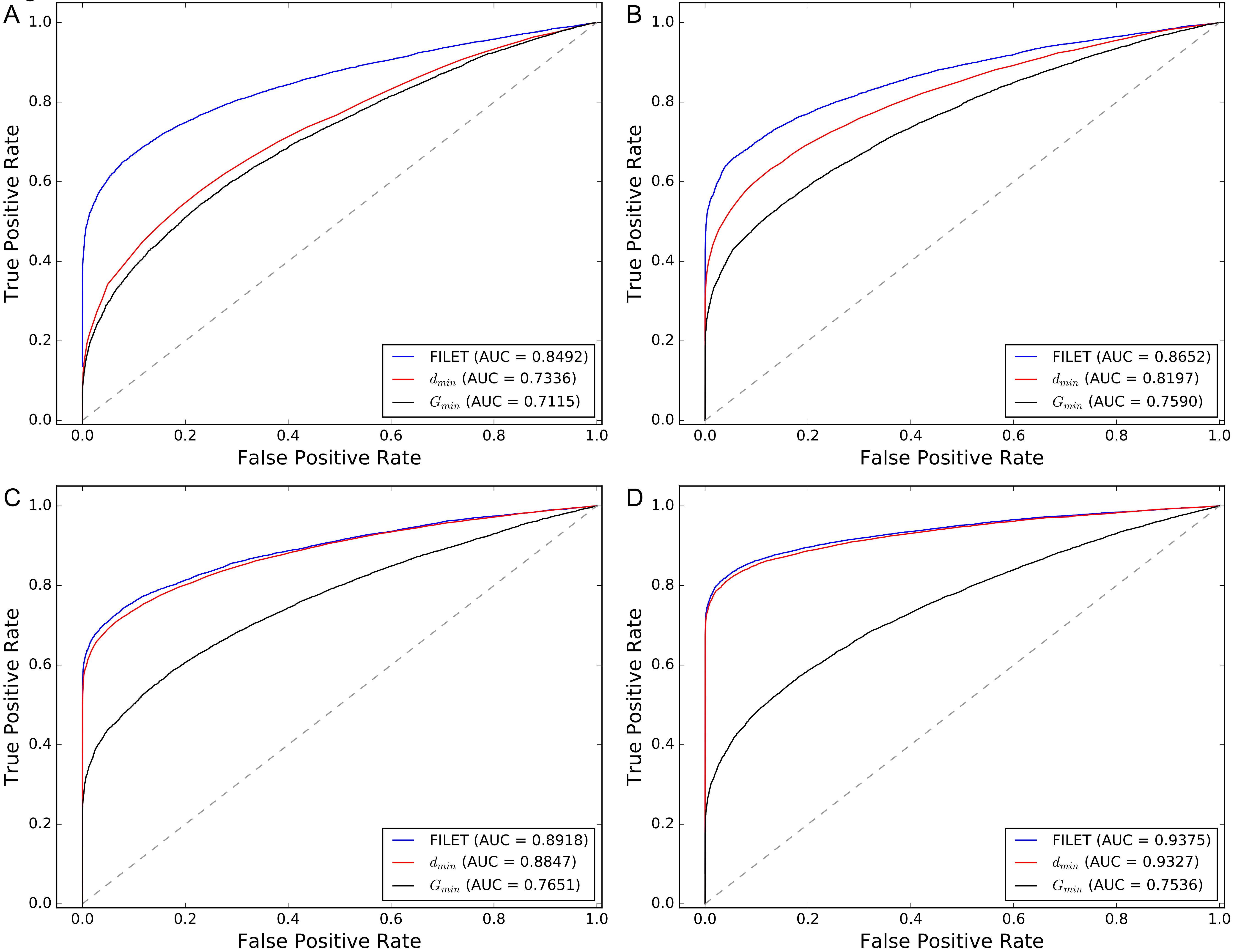

### S6_Fig.pdf

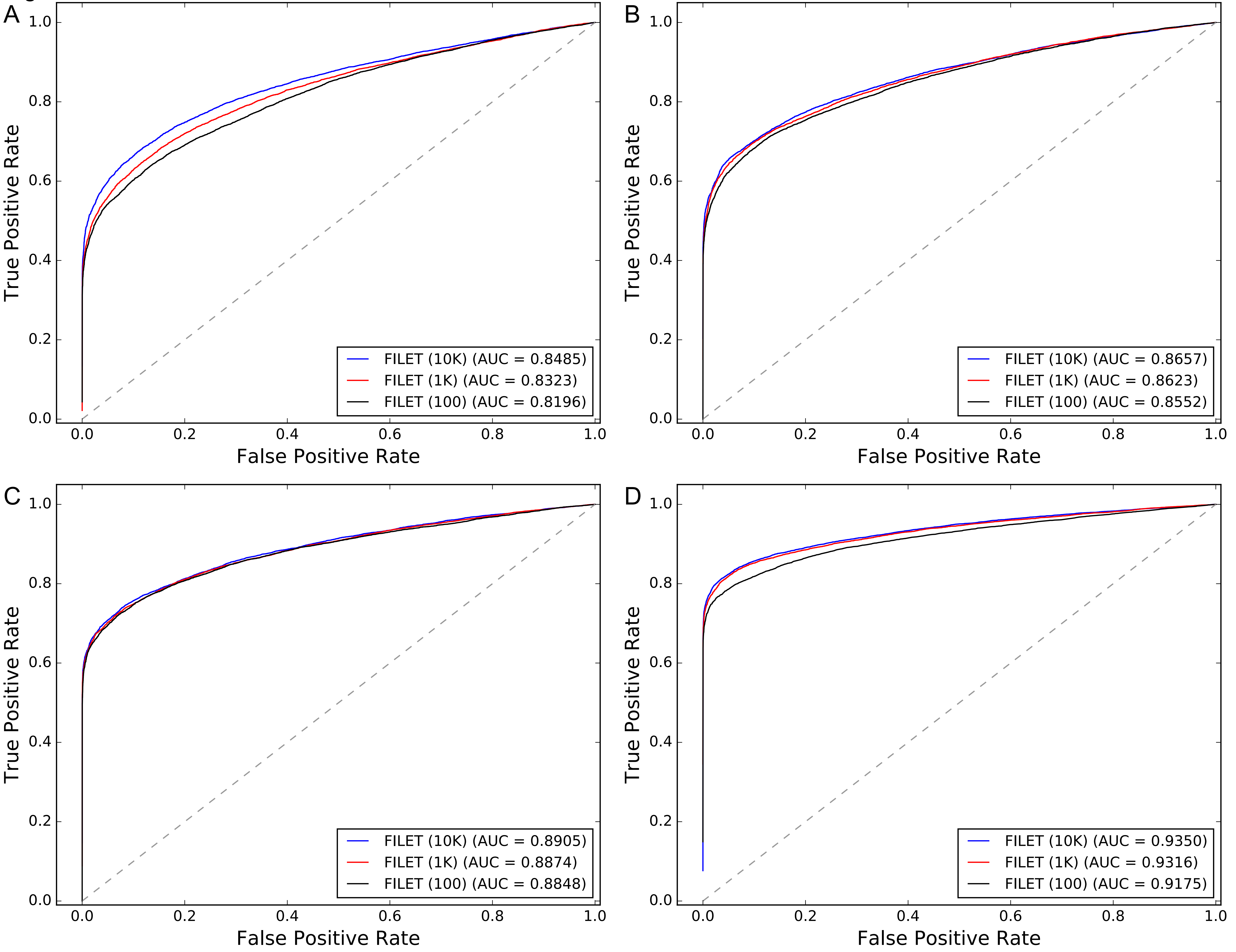

### S7_Fig.pdf

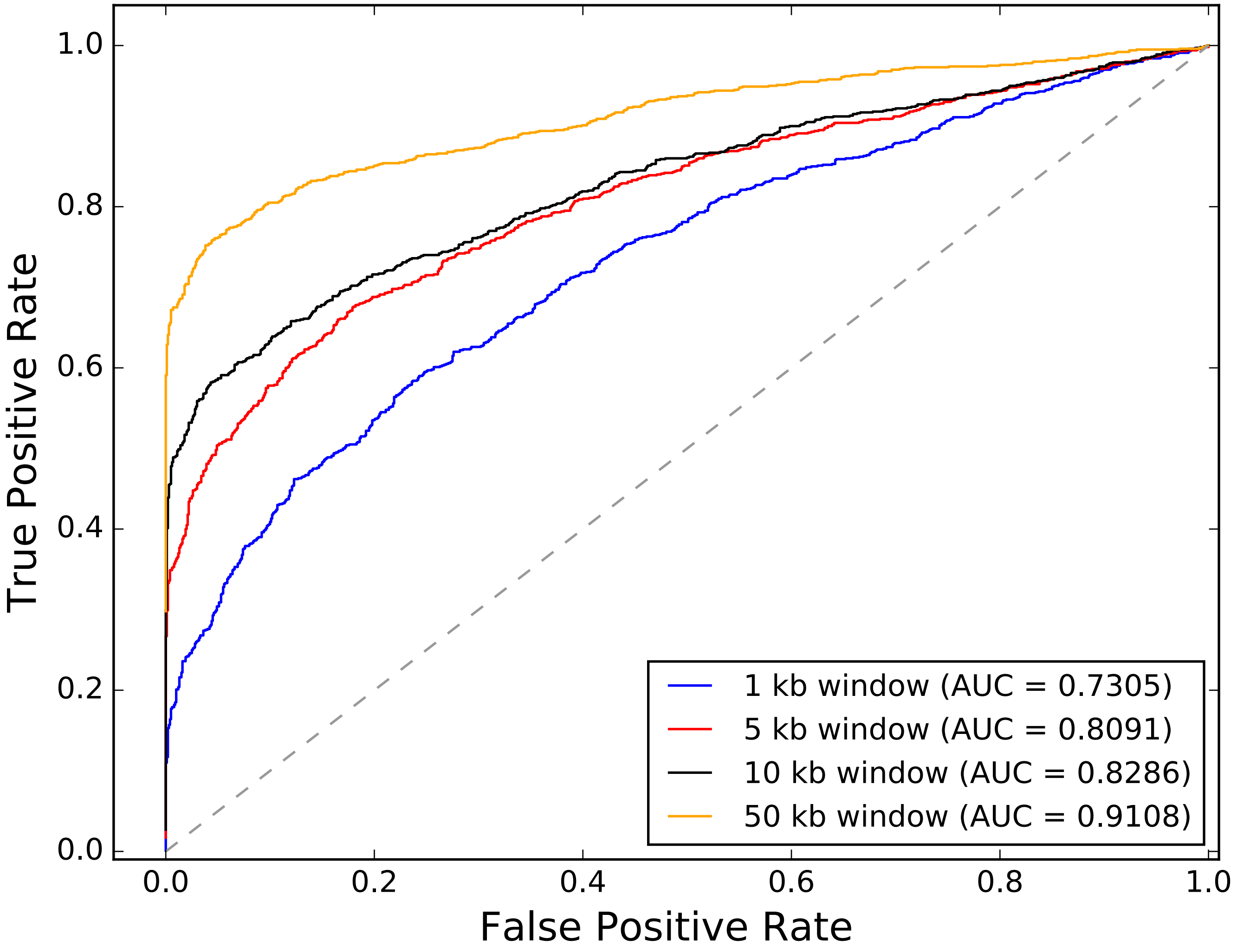

### S8_Fig.pdf

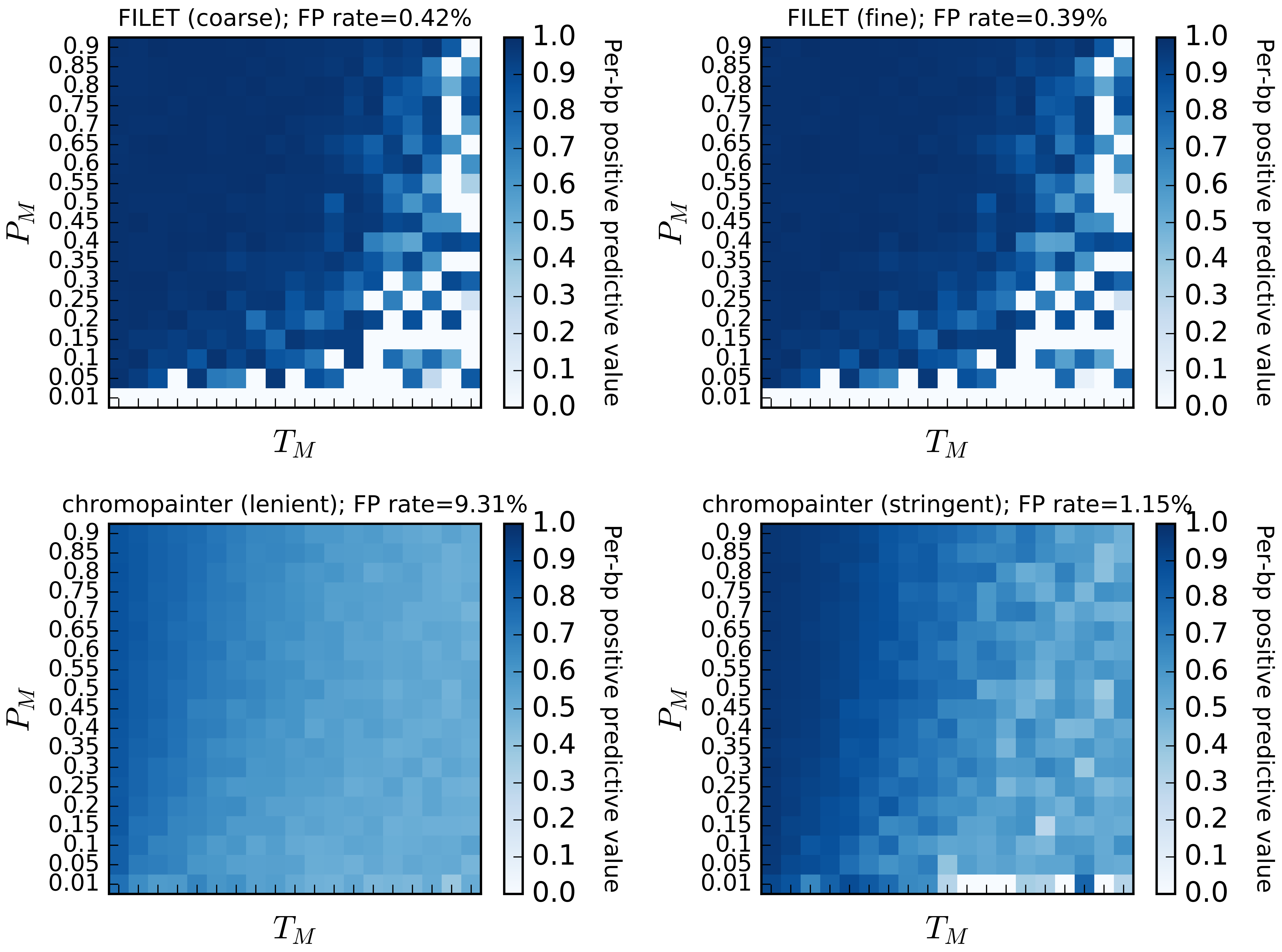

### S9_Fig.pdf

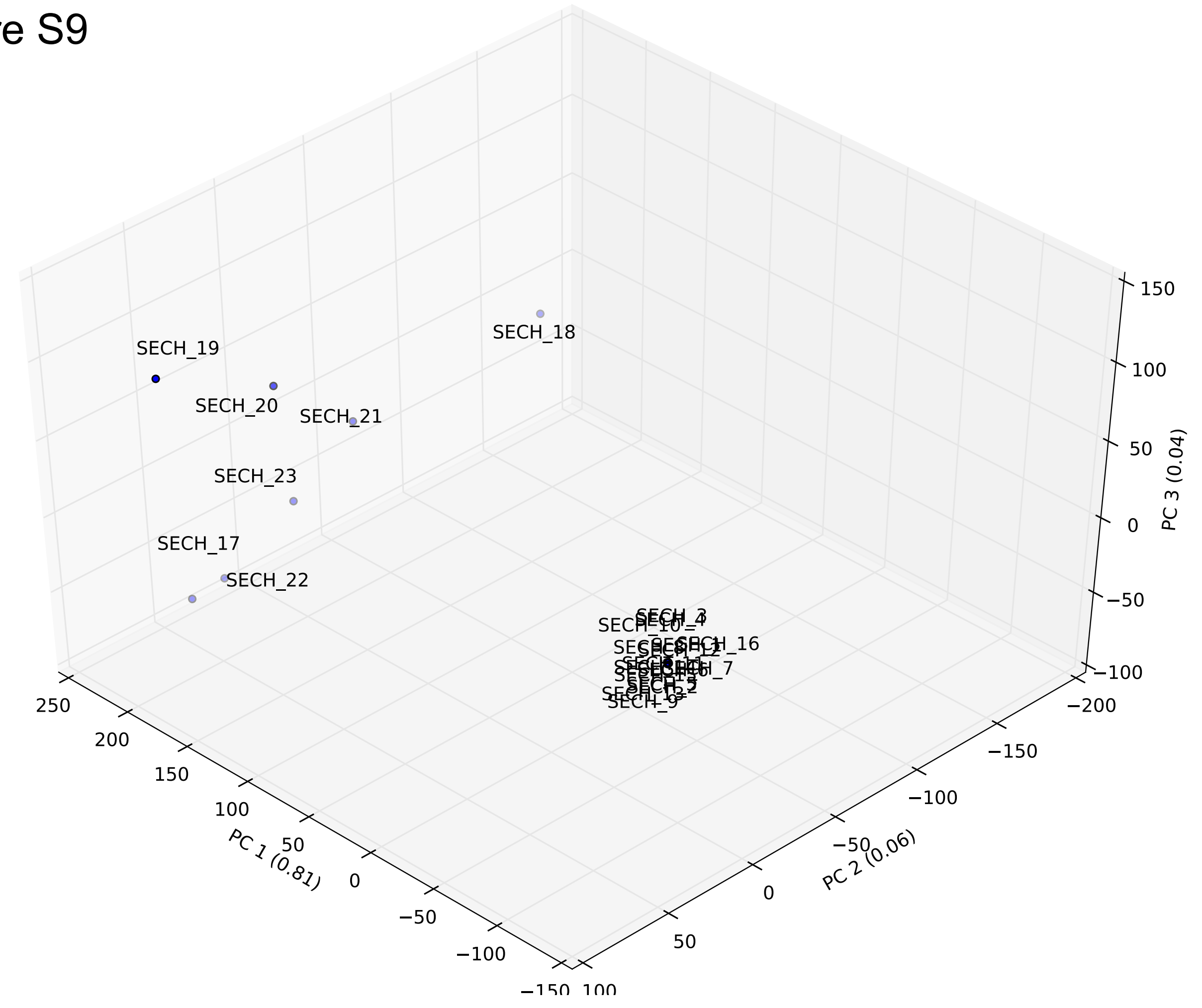

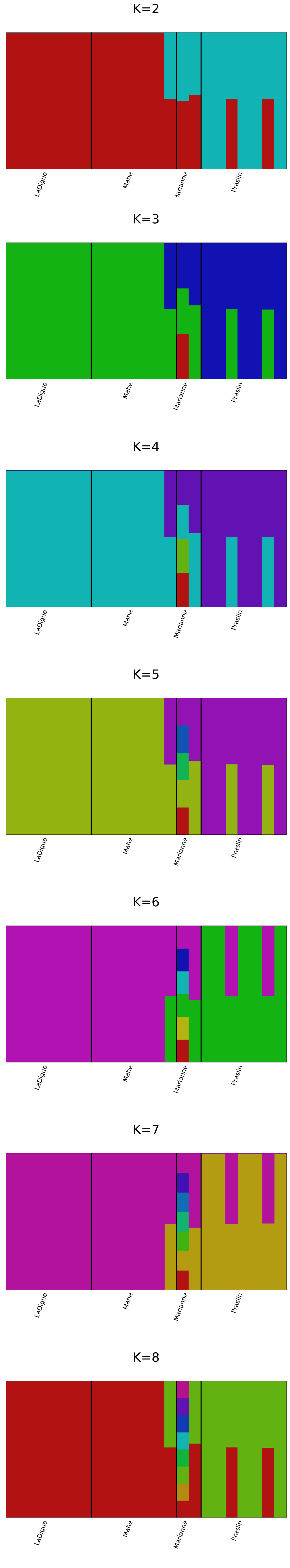

### S10_Fig.pdf

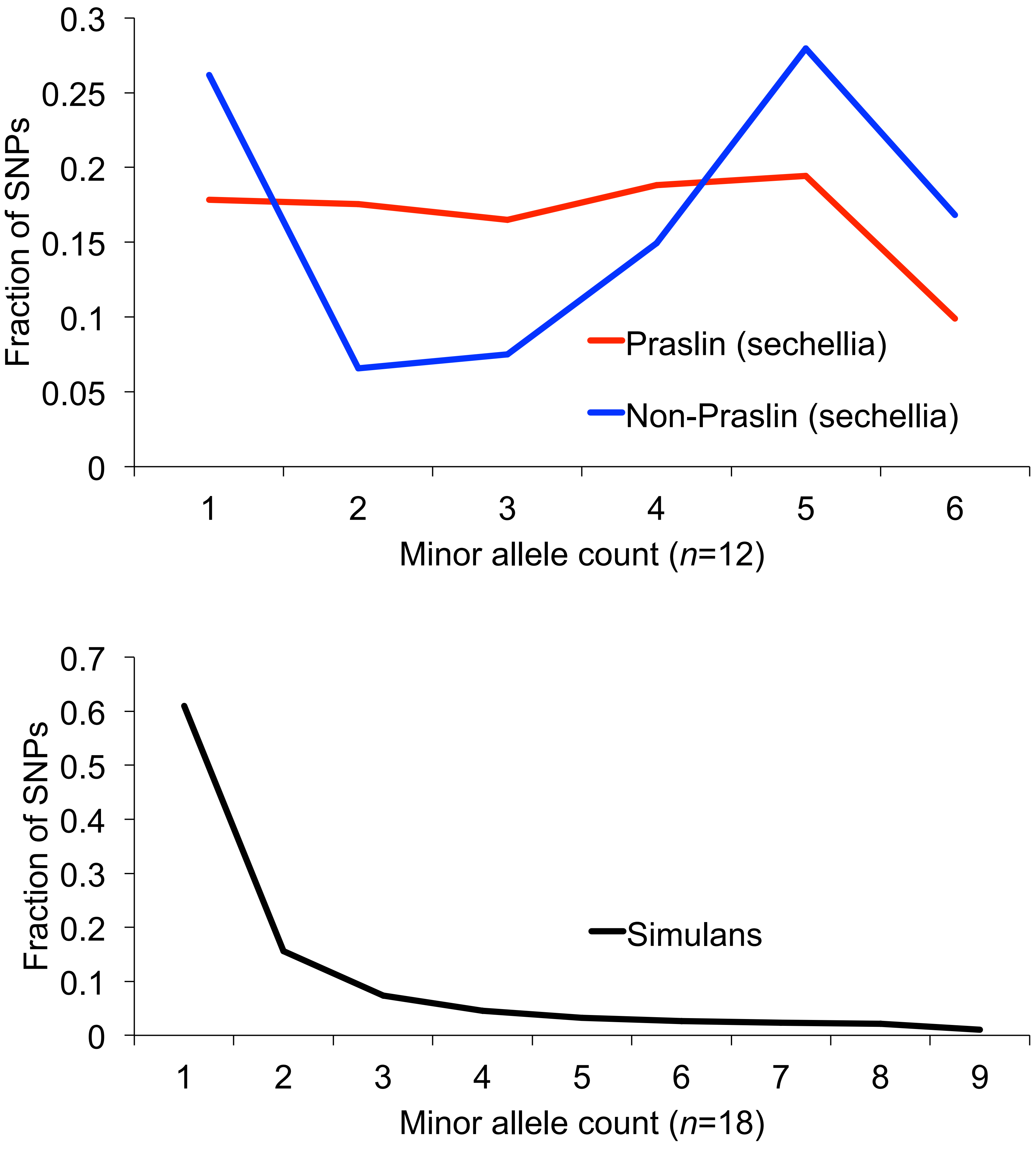

### S11_Fig.pdf

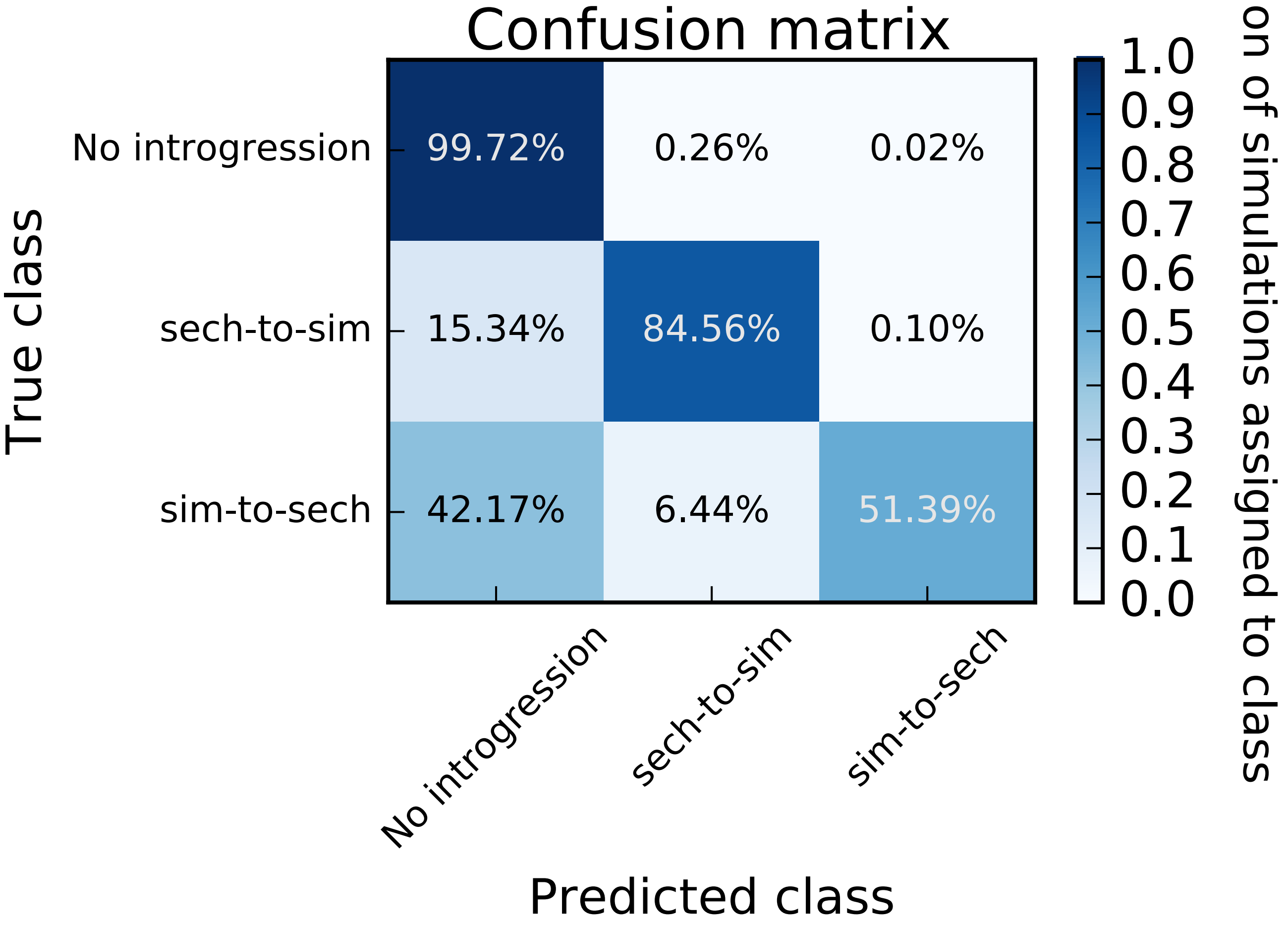
